# The chloroplast ionome shines new light on organellar Fe homeostasis

**DOI:** 10.1101/2025.07.28.667122

**Authors:** Lorenz J. Holzner, Lucia Östergaard Frank, Susanne Mühlbauer, Anna Müller, Laura Schröder, Rachael A. DeTar, Katrin Philippar, David Mendoza-Cózatl, Ute Krämer, Thomas Nägele, Bettina Bölter, Hans-Henning Kunz

## Abstract

Annually, chloroplasts fix 258 billion tons of CO_2_ through photosynthesis. Photosynthesis and other biochemical pathways require specific amounts of metal ions in the organelle. Transport proteins in the plastid inner envelope maintain the organellar ion homeostasis. Despite substantial progress over the last decades, many genes encoding for plastid ion channels and ion carriers or their regulators remain unknown. To fill this knowledge gap, detailed information on the elemental composition of chloroplasts i.e., a plastid ionome, is needed. This will allow to compare mutants of transporter candidates with wild-types.

Here, we provide quantitative descriptions of chloroplast ionomes from *Arabidopsis thaliana,* the metal hyperaccumulator *Arabidopsis halleri*, *Pisum sativum*, and *Nicotiana benthamiana* and analyze similarities and distinctions. Using *A. thaliana,* we show that plastid ionomes can be genetically manipulated. Chloroplasts of *oligopeptide transporter3* (*opt3*)-deficient mutants contain 14-fold more iron, which they deposit into stromal FERRITIN. The removal of FERRITIN in *opt3* mutants leads to a substantial decrease in plastid and leaf iron pointing to important signaling linked to the chloroplast ionome. Our study reveals that chloroplasts can be turned into large iron storages. Since crop biofortification to fight hidden hunger has become a global mission, this research provides groundwork to reach this goal.

## Introduction

Plants rely on specific mineral nutrients to perform basic physiological processes such as pH- and osmo-regulation or enzymatic reactions. The mineral nutrient and trace element composition of an organism – in this case a plant – is called the ionome (Salt et al., 2008). It varies between cell types and growth conditions (Giehl et al., 2023). Root, leaf, and seed ionomes from the model plant *Arabidopsis thaliana* grown under standard conditions were established by Salt and colleagues (Lahner et al., 2003; Salt, 2004; Baxter et al., 2008). Their work coincided with the release of the Arabidopsis genome and the emergence of mutant collections. Subsequently, plant mutants could be evaluated based on their ionome relative to wild-type (Wt) controls. This approach fostered the discovery of genes encoding metal ion transport proteins, their regulators, and nutrient signaling components. A growing database of mutants with altered ionomes documents numerous breakthroughs but reveals many genetic components that remain unknown (Whitt et al., 2020). Since plants represent the foundation of our food chain, we must unveil all missing components of their mineral homeostasis. Only a deep understanding of this aspect of plant physiology will secure future crop yields, enable sustainable agriculture, and fuel innovations such as biofortification (Palmgren et al., 2008) and phytoremediation (Krämer, 2005).

A current blind spot in understanding plant nutrition is the subcellular distribution of mineral nutrients. The chloroplast, with its high demand for iron (Fe), manganese (Mn), copper (Cu), and zinc (Zn) in the photosynthetic machinery, is of particular interest. Almost all types of plant malnutrition affect chloroplast function and consequently plant productivity (reviewed in Kunz et al. (2024)). Deciphering the genes that govern the organellar ionome will enable strategies to protect plastids during phases of adverse nutrient supply. Therefore, a reference chloroplast ionome is urgently needed to provide a solid foundation for this gene exploration. Although the element composition of spinach and pea plastids was published decades ago (Demmig and Gimmler, 1983; Robinson and Downton, 1984; Schröppel-Meier and Kaiser, 1988), chloroplast elemental data from the genetically accessible species *A. thaliana* remain rare and restricted to a singular, or a small subset of ions at a time (Seigneurin-Berny et al., 2006; Jeong et al., 2008).

In this study, we set out to quantify and genetically challenge the chloroplast ionome of the model plant Arabidopsis. For several reasons we focused on the organelle’s Fe level. Firstly, 79% of the leaf Fe content is proposed to localize in chloroplasts (Terry and Low, 1982). Secondly, leaf FERRITINS (FER1, 3, 4), the main Fe storage proteins under excess conditions, locate within plastids (Ravet et al., 2009; Roschzttardtz et al., 2013). Thirdly, biofortification, i.e. raising Fe levels in specific crops, is of great interest to fight Fe deficiency in the human diet which is referred to as “hidden hunger” (Murgia et al., 2012). Turning leaf plastids into Fe sinks may prove a valuable biofortification strategy to fight hidden hunger.

Although Fe is among the most abundant elements in the earth crust, its bioavailability is low (Morrissey and Guerinot, 2009; Colombo et al., 2014). Arabidopsis and other dicots acidify the rhizosphere to mobilize Fe^3+^ chelates. Subsequently, FERRIC REDUCTION OXIDASE 2 (FRO2) reduces ferric chelates to ferrous iron (Fe^2+^), which is shuttled across the plasma membrane of root cells by IRON REGULATED TRANSPORTER 1 (IRT1) and to a lesser extent by NATURAL RESISTANCE ASSOCIATED MACROPHAGE PROTEIN 1 (NRAMP1) (Vert et al., 2002; Cailliatte et al., 2010).

Fe uptake and homeostasis are tightly regulated. A key player is the transcription factor FIT, which controls FRO2 and IRT1 expression. Despite the discovery of many components of the Fe homeostasis regulatory network in the last years, where exactly the plant Fe status is sensed and which mechanisms govern the Fe signaling network is unclear. One sensor candidate is the E3 ubiquitin ligase BRUTUS (BTS). BTS is proposed to bind Fe via its hemerythrin domains and marks FIT and other transcription factors for degradation (Hindt et al., 2017; Rodríguez-Celma et al., 2019). Furthermore, BTS ubiquitinates a class of short peptides named IRON MAN (IMA) peptides. Low internal Fe levels trigger IMA expression to the point that BTS becomes saturated, preventing the degradation of target transcription factors and promoting Fe uptake (Li et al., 2021).

Internally, Fe can be shuttled between xylem and phloem to supply sink tissues. This requires OLIGOPEPTIDE TRANSPORTER 3 (OPT3), a main player in Fe homeostasis. While null alleles are lethal, hypomorphic *opt3* mutants exhibit leaf necroses and overaccumulate Fe, Mn, Zn, and Cadmium (Cd) in above ground tissues (Stacey et al., 2008; Mendoza-Cózatl et al., 2014; Zhai et al., 2014). High metal uptake in *opt3* is caused by a continuous Fe starvation signal leading to constitutive IRT1 and FRO2 upregulation. Where *opt3* mutants compartmentalize Fe, Mn, and Zn within cells has not been analyzed. Chloroplasts, with their high metal demand, represent a plausible option. Arabidopsis chloroplasts supposedly also employ a reduction-based Fe uptake mechanism (reviewed in López-Millán et al. (2016) and Schmidt et al. (2020)). FERRIC REDUCTION OXIDASE 7 (FRO7), a proposed inner envelope (IE) reductase (Jeong et al., 2008), was proposed to produce Fe^2+^, which is shuttled into chloroplasts by PERMEASE IN CHLOROPLASTS 1 (PIC1) (Duy et al., 2007). While *fro7* null alleles are indistinguishable from Wt plants, *pic1* loss of function mutants exhibit a dwarfish albino phenotype. Another Fe import candidate is MITOFERRIN LIKE 1 (MFL1), a homologue of the mitochondrial Fe transporter MIT1 (Tarantino et al., 2011). However, *mfl1* mutants resemble Wt controls. Only when irrigated with excess Fe, *mfl1* chloroplast Fe drops by 25% (Tarantino et al., 2011).

As the published chloroplast Fe uptake and homeostasis mutants display contrasting phenotypes, research should focus on chloroplast Fe status of respective mutants. Here, we establish the chloroplast ionome at control conditions as a baseline for investigating the canonical Fe import mechanism and other plastid metal transport pathways in the future.

## Results

### Determining the chloroplast ionomes of *A. thaliana* and other dicot species

So far, a basic chloroplast ionome for the model plant *A. thaliana* is missing (Kunz et al., 2024). To overcome present limitations, we determined elements in leaf tissue and isolated chloroplasts from the same batch of 21-day old Wt plants grown on standard soil (fig 1A).

All leaf elements were in a similar range as previously reported (Salt, 2004; Völkner et al., 2024) (fig. 1B, fig. S1D). Regarding photosynthetically relevant transition metals, we obtained almost the same value for Fe (our measurement vs Salt, 2004; 100 vs 101 µg*gDW^-1^) and Zn (70 vs 61 µg*gDW^-1^). Higher levels were obtained for Mn (113 vs 64 µg*gDW^-1^) and Cu (8 vs 1.8 µg*gDW^-1^), while nickel (Ni) was lower and close to the detection limit of our system (0.57 vs 1.4 µg*gDW^-1^). The macronutrients potassium (K) (38,200 vs 46,100 µg*gDW^-1^) and calcium (Ca) (41,025 vs 45,000 µg*gDW^-1^) were slightly lower and phosphorous (P) moderately elevated (10,773 vs 9700 µg*gDW^-1^). We also obtained levels for sulfur (S), chlorine (Cl), and rubidium (Rb) (fig. S1D). Leaf Rb levels were slightly elevated compared to previous values (14 µg*gDW^-1^ vs 10 µg*gDW^-1^) (DeTar et al., 2022).

**Figure 1.**
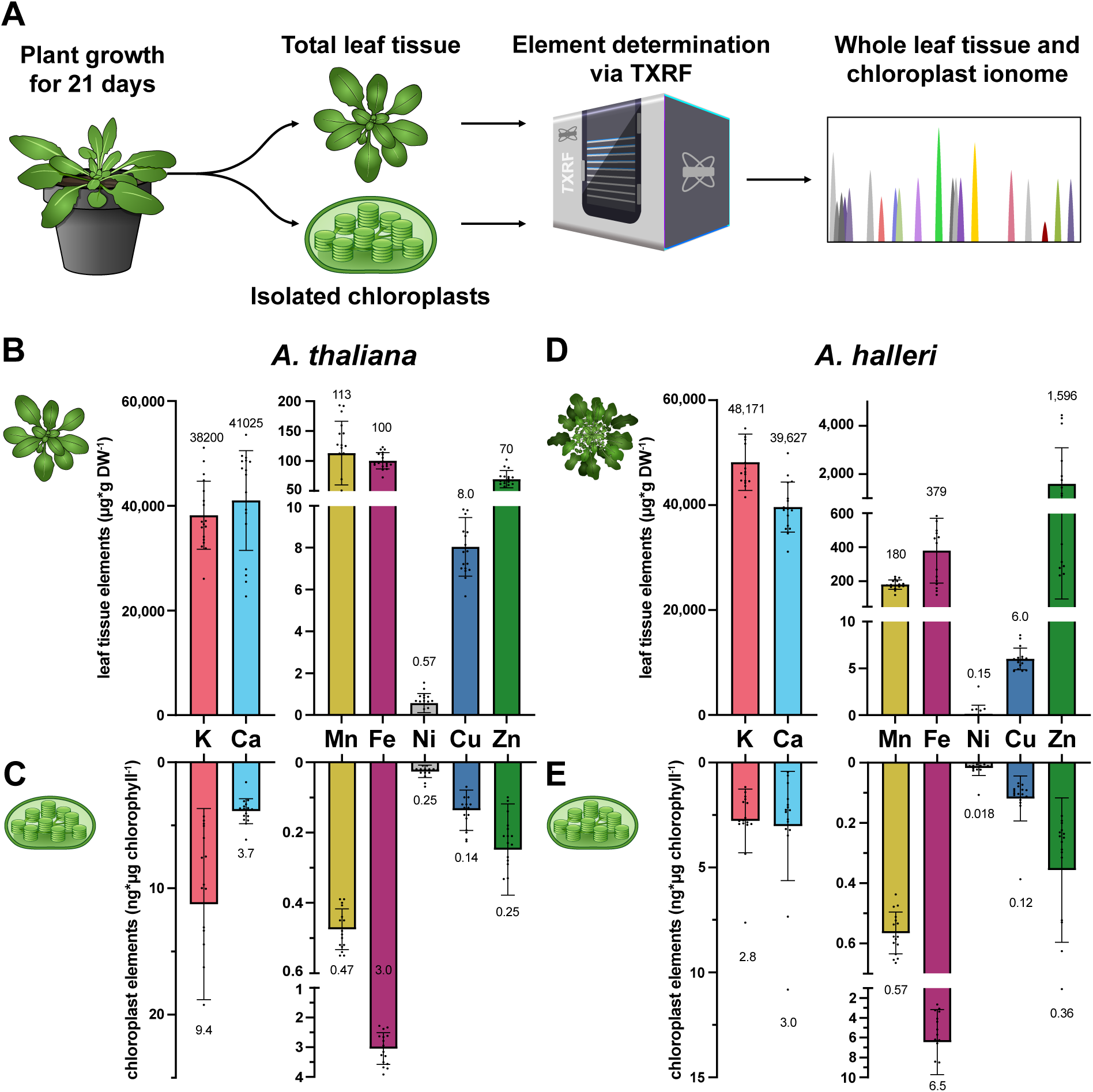
Leaf tissue and chloroplast ionomes of *A. thaliana* and *A. halleri*. **(A)** Schematic outline of sample preparation to determine the leaf tissue and plastid ionome. (**B, D**) Leaf tissue ionomes of *A. thaliana* (*n* = 18) and *A. halleri* (*n* = 16). (**C, E**) Chloroplast ionomes of *A. thaliana* (*n* = 17) and *A. halleri* (*n* = 16). Ionomes including K, Ca, Mn, Fe, Ni, Cu, and Zn were determined using TXRF and normalized to dry biomass (DW; A, C), and chlorophyll (B, D). All data are presented as mean ± SD. Whole-leaf ionomes are from the same batch of leaves that was also used to isolate chloroplasts.

Prior to chloroplast element measurements, we analyzed the quality of isolated chloroplasts from *A. thaliana*, *P. sativum* and *N. benthamiana* via SDS-PAGE, activity measurements of the stromal NADPH malate-dehydrogenase (NADP-MDH), and microscopy (fig. S1A-C). The large subunit of RuBisCO (LSU) was clearly detectable for all species, indicating that only minor amounts of plastid stroma were lost during the isolation procedure (fig. S1A). Activity measurements in sonicated versus non-sonicated chloroplasts showed that isolated chloroplasts from all three species did not display a strong NADP-MDH activity prior to sonication and remained stable for the time period needed to prepare element samples (fig. S1B). Lastly, we analyzed the quality of the chloroplasts by phase-contrast microscopy. Viable chloroplasts display a “halo” (Schulz et al., 2004). In total, 200-300 chloroplasts were evaluated. Based on the presence of a halo, intactness of 72.3% for *A. thaliana*, 77.3% for *P. sativum*, and 71.3 % for *N. benthamiana* was documented (fig. S1C).

Next, we quantified the chloroplast ionome of *A. thaliana* (fig. 1C). We determined mean element levels for the photosynthetic-relevant transition metals: Fe (3.0 ng*µg^-1^ chl.), Mn (0.47 ng*µg^-1^ chl.), Ni (0.25 ng*µg^-1^ chl.), Cu (0.14 ng*µg^-1^ chl.), and Zn (0.25 ng*µg^-1^ chl.). Our Fe values confirmed previous reports by Jeong et al. (2008) (3 ng*µg^-1^ chl). Cu was found in the same range as in measurements by Seigneurin-Berny et al. (2006) (0.12 g*µg^-1^ chl. Cu). Our Zn values were above levels reported by Seigneurin-Berny et al. (2006) (0.1 ng*µg^-1^ chl. Zn). Additionally, we measured Ca (3.7 ng*µg^-1^ chl.) and K (9.4 ng*µg^-1^ chl.).

To expand our insights, we determined leaf tissue and chloroplast ionomes for greenhouse-grown *N. benthamiana* and *P. sativum* (fig. S1D). Compared to *A. thaliana*, we observed significant differences in the leaf tissue ionome regarding Fe, Cu, S, Cl and Ca for *P. sativum*. *N. benthamiana* also varied in Mn, Cu, P, and S contents. As for the plastid ionomes, we detected significant but mostly modest (<3-fold) differences in Mn, Fe, Cu, and Zn for *P. sativum* and for Cu, Zn in *N. benthamiana* chloroplasts. One notable exception was Ca; *N. benthamiana* revealed a 37-fold enrichment compared to *A. thaliana*. Since leaf Ca levels were not significantly altered, the Ca accumulation in *N. benthamiana* plastids may be species-specific.

Subsequently, we addressed a chloroplast ionome from a highly distinct leaf ionome. Therefore, we employed *Arabidopsis halleri*, which exhibits leaf Zn levels more than an order of magnitude higher than in *A. thaliana* and successfully colonizes metal-contaminated mining sites (Krämer, 2010; Stanton et al., 2022). Initially, we compared the leaf ionome of *A. halleri* to *A. thaliana*. Confirming previous studies (Stein et al., 2016; von Bremen-Kühne et al., 2022), we documented similar K and Ca values in *A. halleri.* Ni and Cu were unchanged compared to *A. thaliana*. As expected, Zn, and to a lesser extend Fe and Mn were elevated. Zn was 23-fold elevated in *A. halleri* leaves compared to *A. thaliana* (fig. D). Next, we determined the ionome of isolated chloroplasts from *A. halleri*. Ca, Mn, Ni, and Zn levels were similar, while K was moderately decreased compared to *A. thaliana*. Contrasting the 23-fold increased Zn level in leaf tissue, we observed only a 1.5-fold zinc increase in *A. halleri* plastids (fig. 1E). This indicates that *A. halleri* may actively shield its chloroplasts from Zn overload.

Interestingly, *A. halleri* exhibited a 4-fold increase of Fe in the leaves, which resulted in a 2-fold increase of chloroplast Fe. This bares the question: Does the chloroplast selectively accumulate specific metals such as Fe?

### Chloroplasts can be turned into large Fe storages

Chloroplasts have been suggested as the main sink for leaf Fe, harboring 79% of total cellular Fe (Terry and Low, 1982). To test if the chloroplast ionome can be genetically manipulated, we employed hypomorphic *opt3* mutants. In line with previous studies (Stacey et al., 2008; Mendoza-Cózatl et al., 2014; Zhai et al., 2014), we found dramatically high Fe leaf levels in 21-day old plants with 1499 µg*gDW^-1^ in *opt3-2* and 1795 µg*gDW^-1^ in *opt3-3,* respectively, marking a 15-to 18-fold increase compared to Wild type (Wt) (fig. 2A). Also, Mn and Zn were significantly elevated in *opt3* leaves.

**Figure 2.**
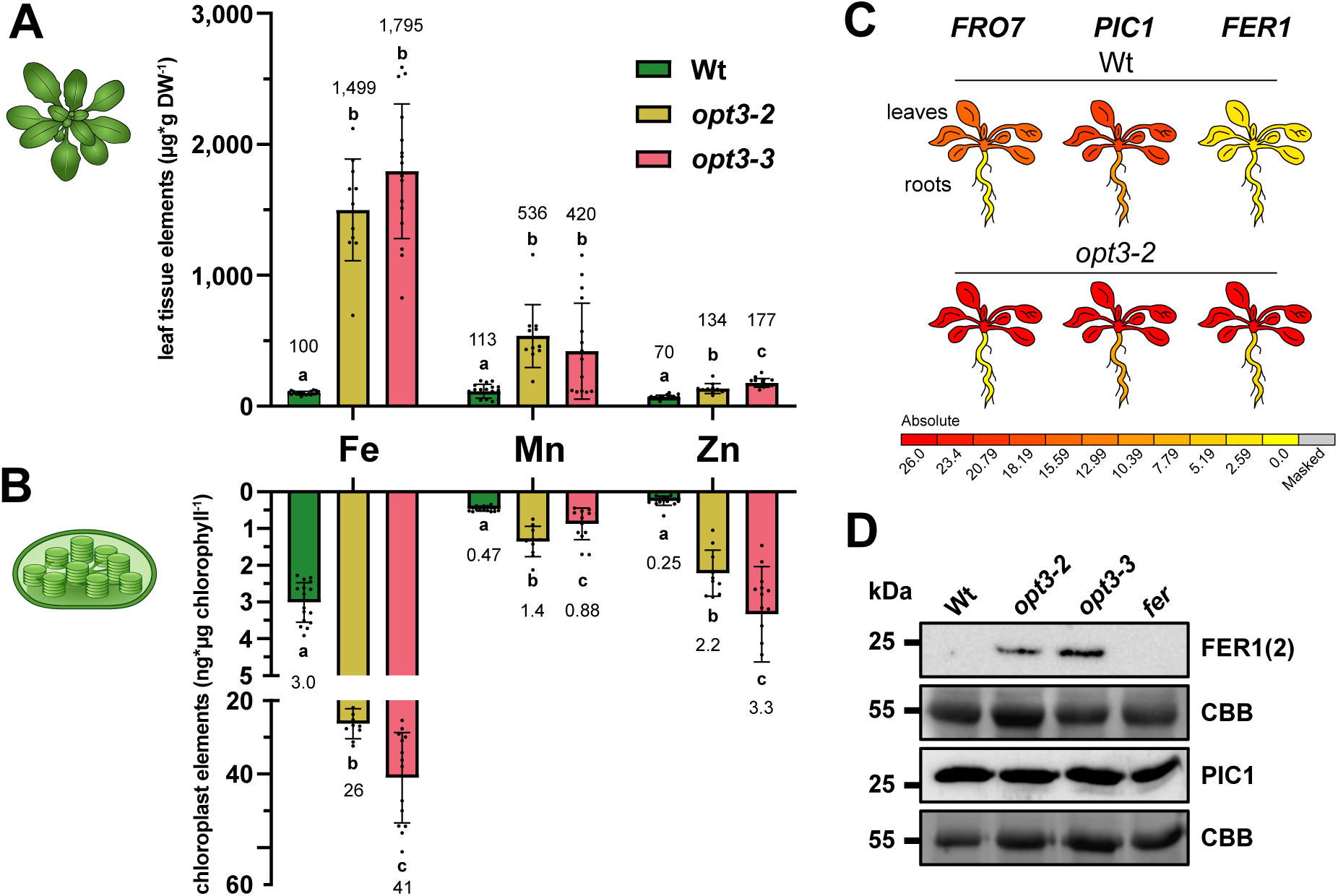
Fe over-accumulated in leaves is stored in chloroplasts. **(A)** Leaf tissue ionomes were determined for the Wt and Fe over-accumulating mutants *opt3-2* (*n* = 11) and *opt3-3* (*n* = 16) using TXRF. Fe, Mn and Zn levels are depicted. (**B**) Chloroplast ionomes were determined from 21-day old Wt, *opt3-*2 and *opt3-3* plants (*n* = 17, 10, and 14 respectively) using TXRF. Samples were normalized to chlorophyll. All data is presented as mean ± SD. Significant differences between Wt and *opt3s* were analyzed using one-way ANOVA *p* < 0.05. (**C**) Differential regulation of FRO7, PIC1, and the plastid Fe storage protein FER1 in *opt3-2* according to McInturf et al. (2021). (**D**) Differential regulation of FER1, OPT3, and PIC1 (Duy et al., 2007) was analyzed by immunoblotting with a specific antiserum against the respective protein.

In parallel, we isolated chloroplasts from the plants used to determine the leaf elements. Whereas Wt chloroplasts had an Fe content of 3 ng*µg^-1^ chl., *opt3-2* revealed 26 and *opt3-3* 41 ng*µg^-1^ chl., respectively, marking a 9-to 14-fold increase of chloroplast Fe compared to the Wt (fig. 2B). This indicates that chloroplasts serve as Fe buffers if leaf levels surpass the physiological range. Following the pattern of Mn and Zn accumulation in the leaf tissue, both metals accumulated mildly in *opt3* chloroplasts.

14-fold elevated plastid Fe should be associated with increased Fe uptake rates, which may be accompanied by transcriptional upregulation of the main chloroplast Fe uptake system. Surprisingly, components of the reduction-based uptake system FRO7 and PIC1 were only slightly upregulated in *opt3-2* (fig. 2C) (McInturf et al., 2021). In contrast, the plastid Fe storage protein ferritin (FER1) was highly upregulated. Similar results were described in an independent study (Chia et al., 2023). For PIC1 and FER1 we obtained antibodies and verified this result on the protein level via immunoblotting. While FER1 was undetectable in Wt, robust expression was observed in both *opt3* mutant alleles (fig. 2D). PIC1 levels in the Wt, both *opt3* alleles, and the *ferritin1*,*3,4* triple loss-of-function mutant (*fer*) were unchanged.

### Loss of ferritins, *FRO7,* and MFL1 in *opt3* results in contrasting phenotypes

As *opt3* chloroplasts display significantly elevated Fe levels, Fe transport components should be highly active. Consequentially, defects in Fe uptake should have a noticeable impact on the chloroplast ionome and on plant performance. Therefore, we tested the contribution of canonical Fe transport components by determining chloroplast Fe levels in *opt3-3* (simply *opt3* from here on) and higher-order mutants. As candidates, we chose the following: a) chloroplast ferritins, as their binding capacity may impact Fe gradients and hence import rates (Laulhere and Briat, 1993; Ravet et al., 2009), b) FRO7, which was previously described as the plastid ferric (Fe^3+^) chelate reductase that generates the transport substrate Fe^2+^ (Jeong et al., 2008), and c) MITOFERRIN LIKE 1 (MFL1), a proposed additional plastid inner envelope (IE) Fe^2+^ transporter (Tarantino et al., 2011). All respective mutant alleles had no distinguishable phenotype from wild-type plants at control growth conditions. T-DNA insertions in all single and higher-order mutants were confirmed by PCR (fig. S2A, B). *pic1* mutants, which have extreme albino phenotypes and are barely viable (Duy et al., 2007), were not considered for our study.

Comparing Wt, *fer* and *fro7* mutants, we observed no significant differences in plant size, the maximal photochemical quantum yield of photosystem II (PSII) *F*_v_/*F*_m_, a proxy for PSII damage (fig. 3A), fresh weight (fig. 3B, fig. S2D), or chlorophyll levels in the leaf tissue (fig. 3C, fig. S2E). As described previously (Chia et al., 2023), *opt3* was significantly smaller and displayed reduced fresh weight. *F*_v_/*F*_m_ was not reduced compared to the Wt. Loss of ferritins in *opt3-3* (*opt3fer*) resulted in further plant size and fresh weight reductions (fig. 3A, B). The characteristic necrotic spots, emerging in older *opt3* leaves (Stacey et al., 2008), were also visible in *opt3fer.* Besides minor changes, the photosynthetic performance of *opt3fer* was largely similar to *opt3* (fig. 3D). Also, major photosynthetic complexes were intact in *opt3fer* (fig. S2F). In contrast to *opt3fer*, the loss of *FRO7* in *opt3* did not affect size or fresh weight in *opt3fro7* mutants (fig. 3A, B, fig. S2C, D).

**Figure 3.**
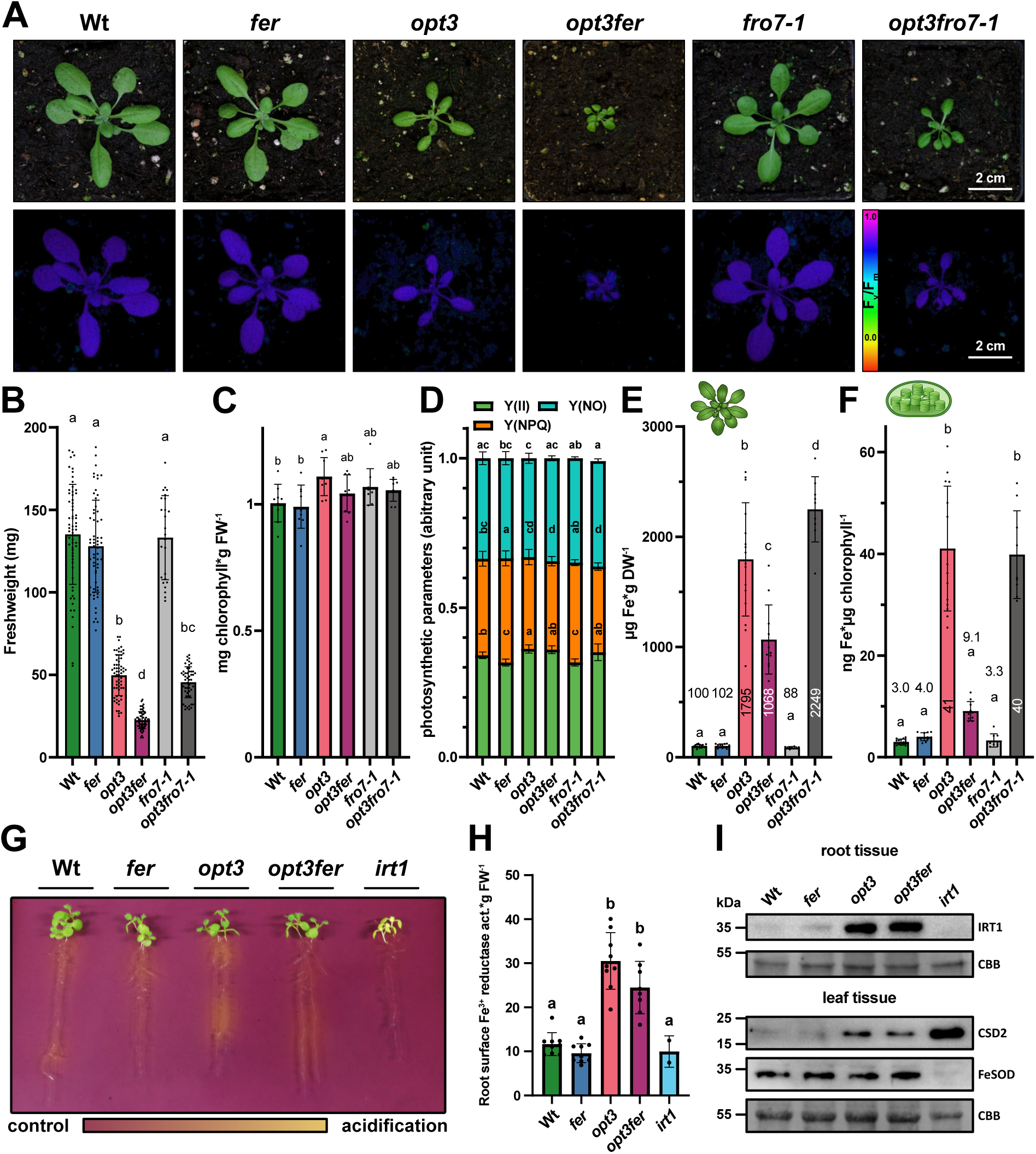
Loss of FRO7 and FER1 in *opt3* display contradictory phenotypes. (A) Phenotype (visual and PAM) of Wt, *fer*, *opt3*, *fro7-1* and higher order mutants at 21 days. (**B**) Fresh weight of mutant lines harvested at 21 days. (**C**) Leaf chlorophyll of mutant lines harvested at 21 days. (**D**) Photosynthetic performance of mutant lines. Y(II), Y(NPQ), and Y(NO) was determined via PAM. (**E**) Leaf tissue Fe levels were measured for Wt (*n* = 18), *fer* (*n* = 12), *opt3* (*n* = 16), *opt3fer (n* = 11), *fro7-1* (*n* = 9), and *opt3fro7-1* (*n* = 9) using TXRF. (**F**) Chloroplast ionomes of Wt (*n* = 17), *fer* (*n* = 11), *opt3* (*n* = 14), *opt3fer* (*n* = 10), *fro7-1* (*n* = 9), and *opt3fro7-1* (*n* = 9) were determined using TXRF. Data of B-F is presented as mean ±SD. Significant differences between lines were analyzed using one-way ANOVA (*p* < 0.05). (**G-H**) 3 components of the root Fe uptake mechanism were analyzed for Wt, *fer*, *opt3*, *opt3fer*, and *irt1* as a control. (**G**) Root acidification was determined by the color change of the pH indicator bromocresol blue from red to yellow. (**H**) Activity of the root surface ferric Fe reductase was measured by Fe^3+^ reduction using Fe^3+^ bathophenanthroline disulfonate (BPDS) an indicator. Data is presented as mean ± SD. Significant differences between lines were analyzed using one-way ANOVA (*p* < 0.05). (**I**) The expression level of IRT1, FeSOD, and CSD2 was analyzed by immunoblotting with a specific antiserum against the respective protein.

Next up, we determined the leaf and chloroplast ionomes for this mutant panel. *opt3fer* mutants revealed a significant drop in leaf Fe by 41% compared to *opt3* (fig. 3E). Mn and Zn levels were also reduced in *opt3fer* compared to *opt3* (fig. S3). The loss of stromal ferritins in *opt3* had a similar strong impact on the chloroplast ionome (fig. 3F). Fe levels in *opt3fer* (9.1 ng*µg^-1^ chl.) were significantly (77%) below *opt3* (41 ng*µg^-1^ chl.) but 3-fold above Wt (3 ng*µg^-1^ chl.). Taken together, this indicates that a lack of chloroplast Fe storage led to lower Fe leaf uptake.

To probe Fe uptake into the root, we initially determined rhizosphere acidification (fig. 3G). Wt and *fer* acidified to a similar extent. In contrast, *opt3* displayed a pronounced rhizosphere acidification, which was attenuated in *opt3fer* and absent in *irt1* plants. Subsequently, root Fe^3+^ reductase activity was quantified as described by (Grillet et al., 2018). Reductase activity was significantly higher in *opt3* and *opt3fer* than in Wt and *fer* genotypes. Reductase activity in *opt3fer* was by trend reduced comparing to *opt3* (≈ 15%) (fig. 3H). Immunoblotting suggested similar levels of the main Fe importer IRT1 in *opt3* and *opt3fer* roots i.e., both genotypes revealed an upregulation of IRT1 protein levels compared to Wt and *fer* plants (fig 3I). In leaves, we documented an upregulation of the chloroplast Fe deficiency marker (CSD2) (Waters et al., 2012) in *opt3* and *opt3fer* whereas no changes were found for Fe superoxide dismutase (FeSOD). Interestingly, in *irt1* mutants FeSOD was undetectable.

In contrast to ferritins, loss of *FRO7* in *opt3fro7-1* and *opt3fro7-2* double mutants did not affect plant growth, fresh weight, *F*_v_/*F*_m_, and photosynthetic performance differently from *opt3* single mutants (fig 3. A, B, D). Both *opt3fro7* alleles showed a 25% increase in leaf Fe compared to the *opt3* mutants (fig. 3E, fig. S3). However, the plastid ionome remained indistinguishable from *opt3* (fig. 3F, fig. S3).

Since our observations indicate that the main chloroplast Fe uptake route may not require FRO7, we designed stable FRO7- and FER1-YFP fusion lines and performed *in vivo* protein localization studies (fig. S4A). Confirming earlier studies (Roschzttardtz et al., 2013), FER1-YFP localized in discrete spots, which perfectly overlaid with the chlorophyll fluorescence signal of chloroplasts. FRO7-YFP resulted in a different pattern. Instead of a ring-like localization surrounding the chlorophyll signal, as expected for a plastid IE protein, FRO7-YFP rather displayed a network-like structure, clearly outside chloroplasts (fig. S4A).

We also localized the closest FRO7 homologue *FRO6.* Overall FRO6 and FRO7 are almost identical (91.2 % amino acid sequence), especially in the N-terminal region, which harbors the putative plastid transit peptide necessary for organellar import (fig. S4C). Jeong et al. (2008) previously reported FRO6 localization to the plasma membrane. As depicted in fig. S4A, we also found FRO6-YFP to localize in similar spots as FRO7-YFP. Microscopic localizations were confirmed by immunoblotting (fig. S4B). While FER1-YFP was present in total protein extracts and isolated chloroplasts, FRO7-YFP was detectable in the total protein extract but not in isolated chloroplasts.

MFL1 was proposed to act in Fe import into plastids (Tarantino et al., 2011). We observed no significant differences in biomass or leaf chlorophyll levels (fig. S2 C, D) for *mfl1* single and *opt3mfl1* higher-order mutants from Wt or *opt3-3,* respectively. While leaf Fe levels for *opt3mfl1* lines were slightly elevated, plastid Fe was reduced by 20-25% compared to *opt3* (fig. S3). However, these changes were non-significant.

### Transcriptomics of *opt3fer* reveals restructuring of the Fe starvation signal

As ferritins and the capacity to store Fe in plastids seem to affect plant Fe homeostasis and uptake in *opt3fer*, we were interested to gain a broad overview of transcriptional reprogramming in leaves of Wt*, fer*, *opt3,* and *opt3fer* using quantitative RNA sequencing.

Initially, all transcripts were used in a cluster analysis to test how similar mutants are based on their transcriptome (fig. 4A). The analysis showed that *fer* and Wt controls are nearly identical i.e., fall into one common cluster. *opt3* and *opt3fer* formed a separate cluster and further separated into genotype-specific sub-clusters. In line with their Wt-like appearance, *fer* plants had only 14 significant DEGs (adj. *p*-value < 0.05; log_2_FC ≥ 0.1; *n* = 4), including the three disrupted ferritin isoforms. In contrast, transcriptomes of *opt3* and *opt3fer* were highly distinct from Wt. We found 5527 significantly up- and 5291 downregulated transcripts in *opt3* versus Wt (fig. 4B, tab. S2). In-depth comparison with two previous *opt3* transcriptome studies (*opt3-2* (McInturf et al. (2021) and *opt3-3* Chia et al. (2023)), revealed that 74% of DEGs (*p* < 0.05, log_2_FC ≥ 0.1) determined in *opt3-2* by McInturf et al. (2021) were also significantly differentially expressed in our dataset (fig. S5A). The majority of these transcripts are similarly up- or downregulated (fig S5D). Comparing our dataset to the two *opt3-3* datasets (old vs. young leaf tissue) determined by Chia et al. (2023), we found that 60% of transcripts in old and 66% in young leaf tissue were also detected in our dataset (fig. S5B). A similar observation was made by comparing the Chia et al. (2023) (*opt3-3* old and young leaves) to the *opt3-2* transcriptome (McInturf et al., 2021) (fig. S5C).

**Figure 4.**
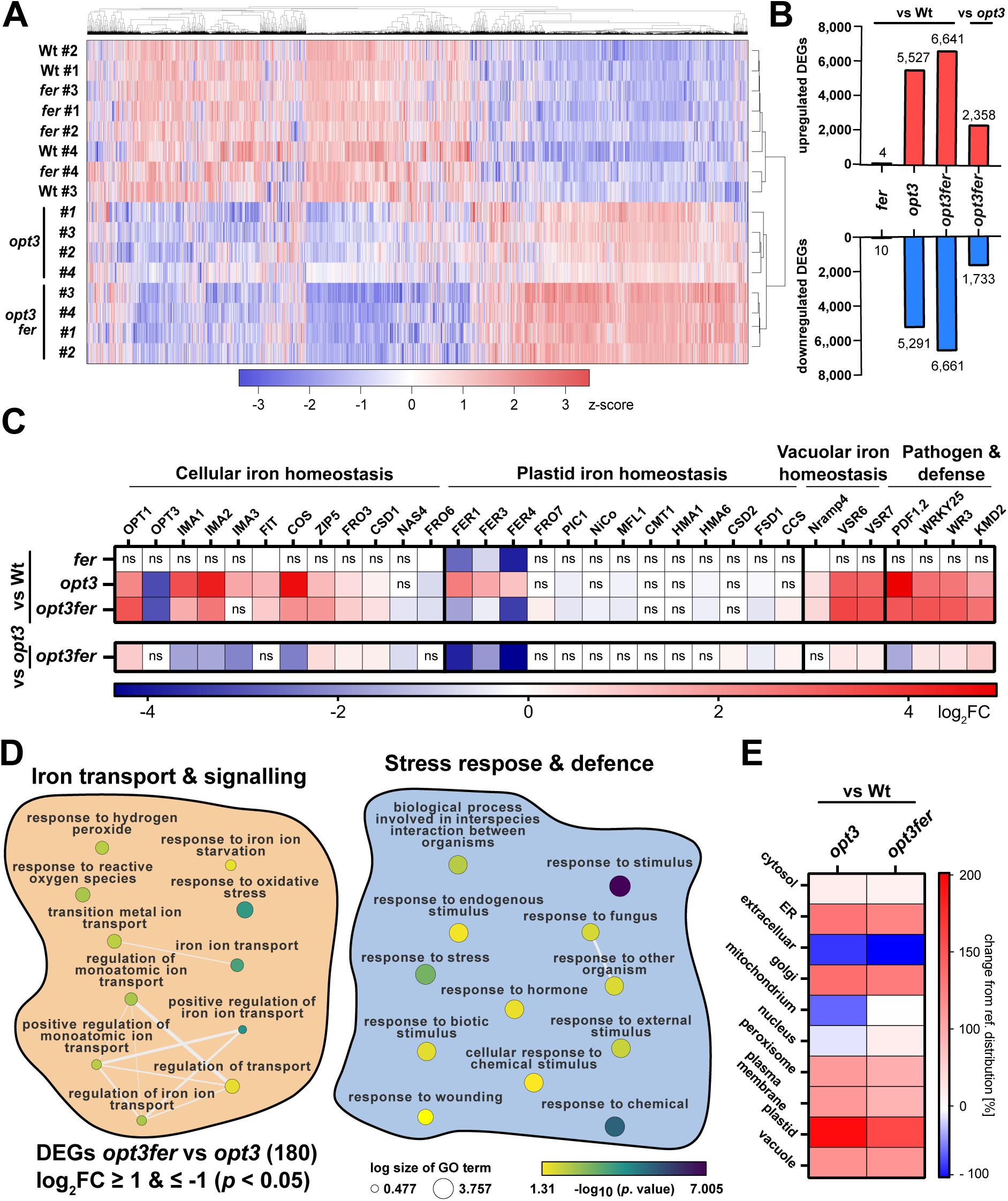
Loss of plastid ferritins in *opt3* influences Fe signaling. **(A)** Total leaf tissue RNA was sequenced and analyzed for differentially expressed genes (DEGs). Normalized counts for all genes of the 4 replicates analyzed (#1, #2, #3, #4) were clustered for similarly expressed genes using heatmap2 on the galaxy platform (Community, 2024). Normalized counts are depicted as z-score per gene. (**B**) DEGs were analyzed for significant DEGs (*p* < 0.05) with a minimum log_2_FC of ± 0.1. (**C**) Significant DEGs were analyzed for ferrome genes defined by McInturf et al. (2021). Additionally, proposed plastid Fe import and homeostasis candidates e.g., PIC1, FRO7, and MFL1 were included. Ferrome genes were grouped into clusters according to their physiological function. (**D**) Strong significant DEGs (log_2_FC **≥** ± 1, *p* < 0.05) of *opt3fer* vs *opt3* were analyzed for gene ontology (GO) terms. The obtained GO terms can be grouped in two main categories: 1. Fe transport and signaling and 2. stress response and pathogen defense. (**E**) Significant DEGs (log_2_FC ± 0.1, *p* < 0.05) were analyzed for subcellular localization using Suba5 (Heazlewood et al., 2006). Data is depicted as relative change to the internal standard distribution of transcripts on Suba5.

For a more specific perspective, we honed in on the genes defined as the leaf ferrome (McInturf et al., 2021) (fig. S5F). A selection of relevant leaf ferrome DEGs is given in fig. 4C. Besides the three leaf-expressed ferritin isoforms (*FER1, FER3, FER4*), no leaf ferrome DEGs emerged in *fer* mutants. In *opt3,* all three ferritin isoforms were significantly upregulated. Also IMA peptides, known for their function in Fe starvation (Grillet et al., 2018), were significantly upregulated in *opt3* (*IMA1* 3.38, *IMA2* 4.33, and *IMA3* 1.68 log_2_FC, respectively). *CAN OF SPINACH* (*COS*), a long non-coding RNA involved in Fe deficiency response (Bakirbas and Walker, 2022), was also significantly upregulated (log_2_FC of 4.65) in *opt3*. Additionally, FIT, the central transcription factor of Fe homeostasis, was upregulated in the *opt3* transcriptome. In summary, the majority (79%) of leaf ferrome (McInturf et al., 2021) loci were significantly (*p* < 0.05) upregulated in *opt3*, confirming the previously described constitutive Fe starvation signaling response in the mutant.

Comparisons between *opt3fer* vs Wt and *opt3* brought further insights. 6641 transcripts in *opt3fer1* were up- and 6661 downregulated compared to Wt. More interestingly, 2385 up- and 1733 downregulated DEGs emerged from *opt3fer* vs *opt3* (fig. 4B). As expected, all three leaf ferritin isoforms were significantly downregulated in *opt3fer* vs Wt and *opt3* (fig. 4C). Interestingly, *bona fide* Fe starvation markers *IMA1*,*2*,*3* were less upregulated in *opt3fer* than in *opt3* (*IMA1* -1.79, *IMA2* -1.72, *IMA3* -2.40 log_2_FC). *IMA3* was not even significantly differentially regulated in *opt3fer* compared to Wt. Also, we detected less *COS* lnRNA transcripts in *opt3fer* than in *opt3* (−2.5 log_2_FC). Lastly, plant defensins (e.g., PDF1.2), which respond to pathogen attacks and bind free Fe in the apoplast (Hsiao et al., 2017), were less expressed in *opt3fer*.

Altogether, the transcriptomes indicate that a loss of plastid Fe storage capacity attenuates the Fe deficiency signal caused by the lack of OPT3, explaining the significantly reduced leaf Fe determined in *opt3fer* (fig. 3E). Interestingly, the canonical plastid Fe import components (*FRO7*, *PIC1*, and *NiCo*, etc.) were at best mildly deregulated in either *opt3* or *opt3fer*.

To gain a broader overview into transcriptome changes upon loss of plastid Fe storage capacity in *opt3* backgrounds, we carried out a Gene Ontology (GO) enrichment analysis using the 180 DEGs identified between *opt3fer* versus *opt3* (cutoff ≤ +/-1 log_2_FC). GO enrichment analysis, using gProfiler and Revigo (Supek et al., 2011; Kolberg et al., 2023), separated the obtained GO terms into two main clusters (fig. 4D): 1) Fe transport and signaling, corroborating lower leaf Fe levels and attenuated IMA, COS, and PDF1.2 expression in *opt3fer*; 2) stress response and defense, containing several genes involved in JA (e.g., JASMONIC ACID CARBOXYL METHYLTRANSFERASE (JMT) and TERPENE SYNTHASE 03 (TPS03)) and Auxin (small auxin up-regulated RNAs (SAUR), Auxin/indole-3-acetic acid 6 (Aux/IAA6)) signaling, as well as additional plant defensins. In general, the majority of genes within the cluster: Fe transport and signaling (e.g., ferritins and iron man peptides) were also found in the other main cluster: stress response and defense, underpinning strong links between Fe homeostasis, pathogen defense, and hormone signaling (Kobayashi, 2019; Herlihy et al., 2020; Liu et al., 2020). The stress response and defense cluster contained three more noteworthy loci: ARACIN1 (ARACIN1) and WALL ASSOCIATED KINASE-LIKE 4 (WAKL4) were both strongly upregulated in *opt3fer*. While ARACINs are Brassicaceae-specific peptides with antifungal function (Neukermans et al., 2015), WAKL4 influences Fe level by inactivating the alternative Fe importer NRAMP1 (Yuan et al., 2024). Lastly, FUSCA3 (FUS3), a transcription factor activated during Fe starvation (Schwarz et al., 2020), was strongly suppressed in *opt3fer* (tab. S2).

SUBA5 (Heazlewood et al., 2006) was employed to determine the subcellular localization of DEG products. *opt3* and *opt3fer* were both enriched in plastid and vacuolar DEGs compared to Wt (fig. 4E). However, in *opt3fer* less DEGs were enriched in plastids whereas the enrichment in vacuolar DEGs was similar to *opt3*. This may indicate a shift towards other subcellular Fe storages in the case when plastid ferritins are absent.

### The central vacuole can act as a backup site for cellular Fe storage

An orthogonal approach to compare element contents of different cell compartments is non-aqueous fractionation (NAF). This technique was applied by Terry and Low (1982) to analyze Fe distribution in leaf cells of sugar beet. Three-week-old Wt, *opt3*, and *opt3fer* plants were subjected to NAF. The relative distribution of the marker enzymes used to quantify organellar enrichments did not vary among genotypes (fig. 5A). Therefore, organelle element distributions can be compared by NAF within these genotypes. Subsequently, the six NAF fractions were subjected to element analysis by TXRF. Elements per organelle were calculated as described previously for metabolites (Hernandez et al., 2023).

**Figure 5.**
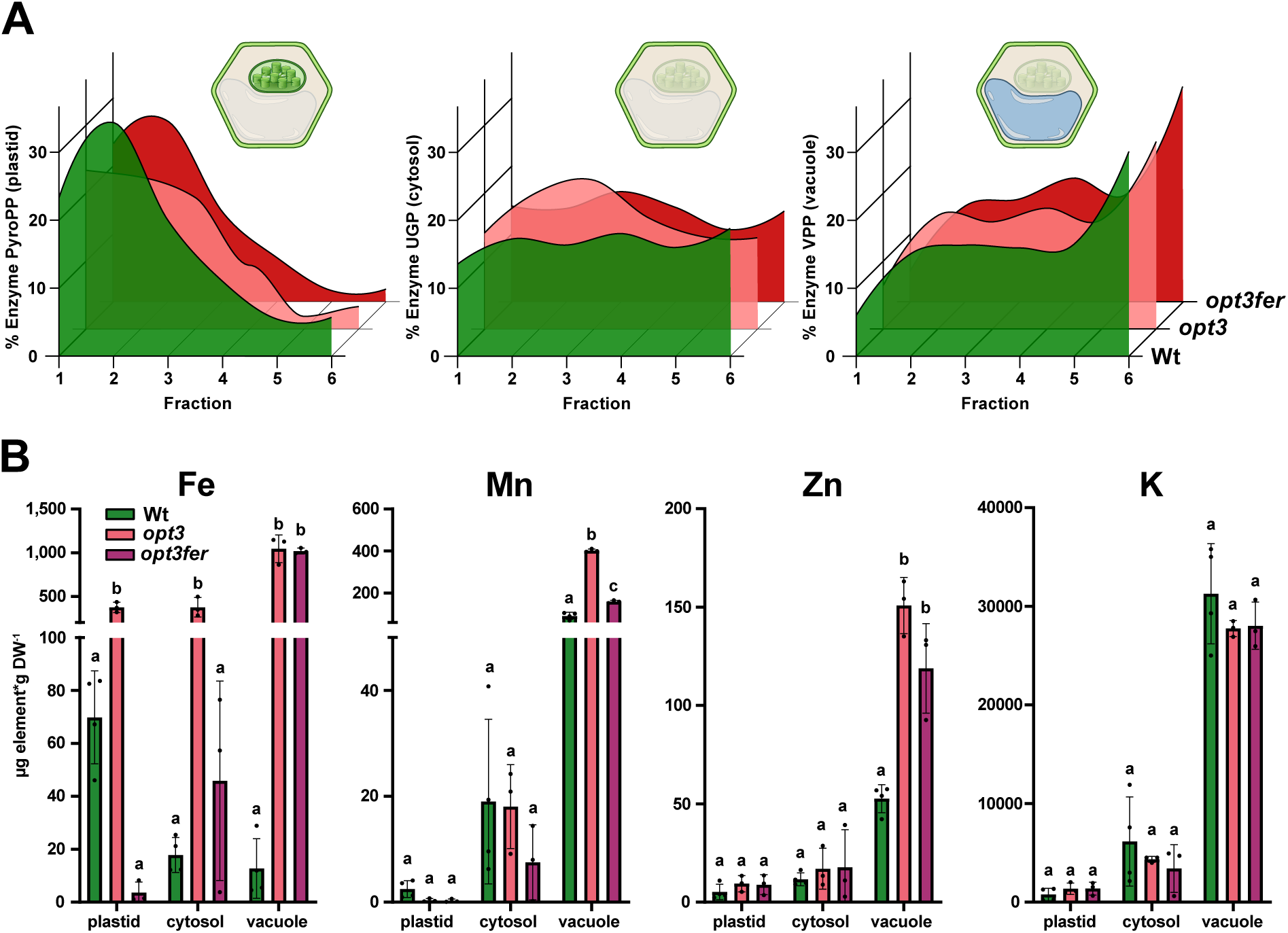
Subcellular distribution of elements. To gain information upon shift in subcellular localization of elements, in particular Fe, upon loss of plastid Fe storage capacity in *opt3fer* we conducted NAF on Wt, *opt3*, and *opt3fer* (**A**) Distribution of the marker enzymes used to trace cellular compartments in the NAF fractions 1-6. No major differences in the distribution of the respective marker enzymes between Wt, *opt3*, and *opt3fer* were obtained. (**B-E**) Fe, manganese, zinc and potassium distribution between organelles of Wt, *opt3*, and *opt3fer*. The relative distribution of the respective element between organelles obtained by NAF was multiplied with the mean dry weight content of the respective element. Data is presented as mean ± SD, *n* = 3-4. Significant differences between Wt, *opt3*, and *opt3fer* within one organelle was analyzed via one-way ANOVA (*p* < 0.05). A full set of elements and relative distribution in % is depicted in fig S6.

Similar to Terry and Low (1982), we found in the Wt 70% of total cellular Fe to be localized to chloroplasts (70 µg Fe*gDW^-1^). 18% were located in the cytosol, and only 12% in the vacuole (fig. S6A, fig. 5B). In *opt3*, we found an increase in plastid iron concentration to 387 µg Fe*gDW^-1^, which equals a 5.5-fold increase in *opt3* comparing to the Wt. In relative distribution this resembles 21% of total cellular Fe content. Further 21% were localized to the cytosol and 58% to the vacuole (388 and 1081 µg Fe*gDW^-1^, respectively). In contrast, for *opt3fer* 95% of total cellular Fe was detected in the vacuole (1036 µg Fe*gDW^-1^). Only 0.3% / 3.6 µg Fe*gDW^-1^, a fraction of Wt levels, was found in *opt3fer* chloroplasts.

Distinct from Fe, the majority of Zn in Wt was found to localize in the vacuole. This was more pronounced in Zn-accumulating lines *opt3* and *opt3fer*. Similar trends were observed for Mn. Lastly, elements not significantly altered among genotypes (fig. S3), were also investigated in NAF fractions. As expected, no genotype-specific differences were observed for K (fig. 5B), Ca, and other elements (fig. S6B).

### Iron starvation response in ferritin-deficient plants

Since in *opt3* ferritins store high amounts of Fe, we were interested how plants deprived of plastid ferritins (*fer* and *opt3fer*) respond to iron starvation. The possibility of iron being released from plastid ferritins was suggested by *in vitro* experiments using pea ferritins (Laulhere and Briat, 1993). However, iron starvation response of *fer* plants has not been analyzed so far.

Wt, *fer*, *opt3*, and *opt3fer* were grown on 1/10 Hoagland media to supplement Fe as desired. An experimental overview is shown in fig. 6A. In brief, plants were germinated on three different media: 1) 0 µM Fe-EDDHA as “Fe free”, 2) 25 µM as “control condition”, and 3) 100 µM as “elevated Fe”. At an age of 10 days, plants were shifted either to fresh media containing the same Fe concentrations serving as control conditions, or to Fe starvation media containing 0 µM Fe-EDDHA. 14 days after the media shift, plants were analyzed for photosynthetic performance and chloroses. Afterwards, rosettes were harvested to quantify the leaf ionome. Determination of the plastid ionome was not possible due to the limited amounts of plant biomass in these experiments.

**Figure 6.**
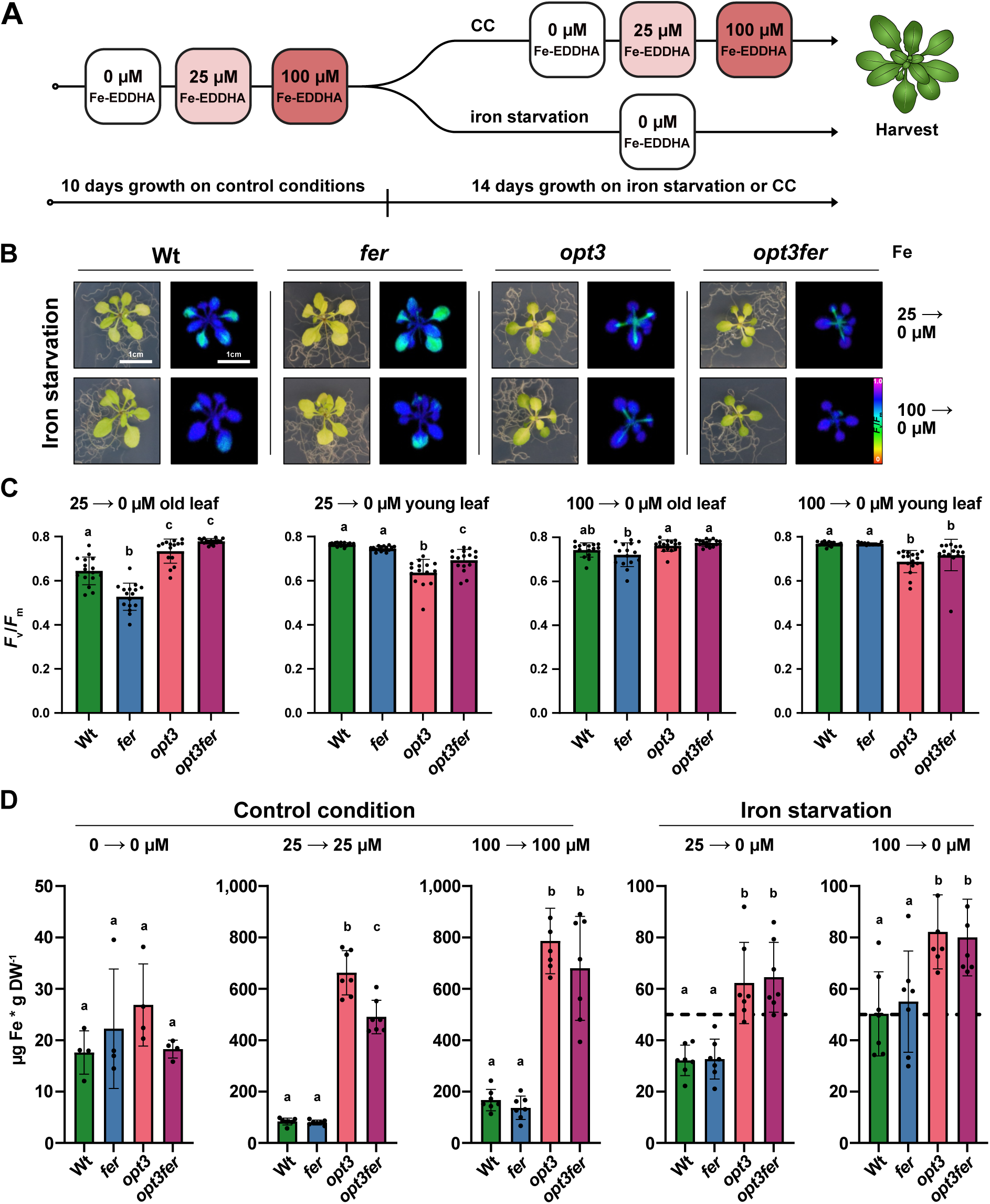
Iron starvation response in ferritin-deficient plants. **(A)** Schematic outline of the Fe starvation assay. Wt, *fer*, *opt3*, and *opt3fer* were cultivated for 10 days at control conditions (0 µM, 25 µM, 100µM). Subsequently seedlings were transferred onto either media containing the same Fe concentration (control condition; CC) or on Fe free media. After 14 days, plants were analyzed by PAM and harvested for element analysis. (**B**) Fe starvation treatment of Wt, *fer*, *opt3*, and *opt3fer*. (**C**) Photosynthetic capacity (*F*_v_/*F*_m_) separately analyzed in old and young leaves resembled the observation displayed in B. Data is presented as mean ± SD, *n* = 16. Significant differences were analyzed via one-way ANOVA. (**D**) Fe levels in plants cultivated on control conditions (0 µM, 25 µM or 100 µM Fe) or after shift from 25 µM/100 µM to Fe starvation. Fe levels were determined via TXRF. Data is presented as mean ± SD, *n* = 4 for 0 µM Fe, *n* = 7 for all other cultivations. Significant differences between Wt, *opt3*, and *opt3fer* within one treatment was analyzed via one-way ANOVA.

All genotypes constantly grown on Fe starvation media displayed a uniform chlorotic phenotype (fig. S7A), reduced *F*_v_/*F*_m_, and Fe levels well-below the defined Fe starvation threshold of 50 µg Fe*gDW^-1^ (fig. 6C) (Jones, 2020). Additionally, we found an increase of Mn and Zn in the Fe-deficient plants due to the broad substrate specificity of IRT1 (Korshunova et al., 1999). Plants grown at 25 and 100 µM Fe-EDDHA throughout the experiment did not exhibit chloroses or decreased photosynthetic capacity and displayed lower Mn and Zn levels. As expected, Fe level increased in all genotypes with rising Fe media contents (fig. S7B). Similar to plants cultivated on soil (fig. 3E, fig. S3), *opt3* contained the highest, *opt3fer* the second highest leaf Fe levels.

We first compared Wt and *fer* shifted from control to iron starvation conditions. Fe-starved plants germinated at high Fe conditions (100 µM Fe-EDDHA) displayed less severe chlorosis, especially in the old leaves, elevated *F*_v_/*F*_m_, and tissue Fe levels compared to the same genotype germinated at lower Fe levels (25 µM Fe-EDDHA) (fig. 6B). If plastid ferritins function as transient Fe storages, which can be remobilized at Fe limiting conditions, *fer* mutants should perform worse than the Wt if shifted from Fe accumulating conditions (100 µM) to Fe starvation. We observed in both Wt and *fer* that older leaves turned chlorotic with concomitant low *F*_v_/*F*_m_ while newly emerging leaves were green with normal *F*_v_/*F*_m_. For plants shifted from 25 µM Fe to 0 µM this drop in *F*_v_/*F*_m_ was significantly stronger in *fer* than in the Wt. Regarding plants cultivated at 100 µM Fe prior shift to 0 µM, no significant difference in *F*_v_/*F*_m_ between Wt and *fer* was observed. Overall, *F*_v_/*F*_m_ was elevated for both Wt and *fer* in this condition (fig. 6C). This indicates that ferritins are not the only Fe release mechanism during Fe starvation conditions which follow Fe surplus periods. We conclude that the positive effect of growing plants on elevated Fe conditions prior to starvation originates from alternative, non-ferritin storages e.g., the vacuole as we have shown in our NAF analysis (fig. 5B), the xylem or apoplastic Fe (apoFe) (Liu et al., 2023).

Next up, we analyzed the Fe starvation tolerance in plants with elevated tissue Fe levels with (*opt3*) and without plastid ferritins (*opt3fer*). In line with a previous study on *opt3* (Stacey et al., 2008), we observed less chlorosis in old leaf tissues of *opt3* and *opt3fer* but pronounced chlorosis in young leaves of *opt3* and *opt3fer* plants (fig. 6B). Consequentially, *F*_v_/*F*_m_ was significantly reduced in young leaf tissues compared to the old leaf tissue for both genotypes (fig. 6C). This effect was less prominent if *opt3fer* plants were cultivated at 100 µM Fe prior Fe starvation. This indicates that forced relocation of cellular Fe into other storages e.g., through the absence of plastid ferritins, seems to provide a slightly larger pool of re-utilizable Fe in *opt3fer*.

## Discussion

To advance understanding about the flux of metal ions into and out of the chloroplast, we set out to quantitatively capture the plastid ionome. Although we focused on the model plant *A. thaliana*, we also analyzed its heavy metal hyperaccumulating sister-species *A. halleri,* and more distant species known for sturdy chloroplasts i.e., *P. sativum* and *N. benthamiana*. In general, we saw similar trends when metal levels were normalized to chlorophyll. Many thylakoid proteins e.g., the photosystems are well-known for their high content of complexed transition metals (Schmidt et al., 2020; Kunz et al., 2024). In addition, also the negatively charged thylakoid membrane itself interacts with divalent cations such as Ca^2+^ (Kaňa and Govindjee, 2016)). This explains why respective elements delivered highly reproducible values across plastid isolations. We conclude that plastid Fe, Zn, Mn, Cu, and Ca values should closely represent the *in vivo* situation. During the isolation process Arabidopsis chloroplasts can briefly open (Schulz et al., 2004). Therefore, some leakage of stroma, where most K resides, may occur. Even though we further optimized our isolation protocol, a slight decrease in the RBCL/LHCII ratio indicative of temporary envelope opening was observed (fig. S1A) and explains the higher variance for K values. We conclude that *in vivo* K in *A. thaliana* chloroplasts may be more abundant. On the contrary, K contents of highly robust pea plastids were slightly below *A. thaliana* with smaller standard deviation. As shown previously, chloroplast K contents can indeed vary among plant species (Robinson and Downton, 1984). For now, we assume plastid K contents in pea and *A. thaliana* are similar. Plastid Ca^2+^ contents in pea and tobacco were 3 and 20-fold higher, respectively, compared to *A. thaliana*. Hence, pea and especially *N. benthamiana* plastids seem to accumulate more Ca.

In general, it seems that plants protect the chloroplast ion homeostasis from metal excess. For instance, plastid Zn contents were fairly stable across all species tested. Even *A. halleri,* which exhibits 20-fold higher leaf Zn level, had no significantly increase in plastid Zn content.

One motivation for this study was to show that genotype-specific plastid ionome analyses can provide valuable mechanistic insights into metal exchange from the organelle and means by plastids to respond to altered nutrient supply. In this regard, we found that *opt3* alleles represent a formidable tool. While it was known that mutant leaves accumulate Fe, Mn, Zn, and Cd (Stacey et al., 2008), it was unknown where the metals are deposited within a cell. Our data show that the mutant uses the chloroplast to sequester the Fe surplus. Up to 14-fold more Fe is shuttled into the organelle than in Wt. This stark difference from the wild type allowed us to revisit some aspects of plastid Fe storage and transport. According to transcriptomics (this study, McInturf et al. (2021), and Chia et al. (2023)) and immunoblots, ferritins, the stromal Fe-binding proteins, are strongly upregulated in *opt3* mutants. When this was prevented by disrupting the three main *FER* loci, resulting quadruple mutants grew significantly slower. Also, the plastid Fe content dropped ≥ 4-fold compared *opt3*. Interestingly, a lack of the reductase FRO7 in *opt3* had no effect on plant growth or plastid iron content. In line with this, we found that neither FRO7 nor its close homolog FRO6 localize to plastids based on fluorescence microscopy. Additionally, publicly available mass-spectrometry data have not detected FRO6/7 in the chloroplast proteome (Heazlewood et al., 2006). Taken together, we conclude that FRO7 likely does not play a major role in the bulk iron uptake mechanism at standard growth conditions. Since no other FRO Fe reductase family member exists in the organelle, several scenarios are conceivable: a) Fe^3+^ uptake is possible (Bughio et al., 1997; Solti et al., 2014) e.g., as Fe^3+^-citrate akin to bacteria (Moraleda-Muñoz et al., 2019) or Fe^3+^-nicotianamine (Seregin and Kozhevnikova, 2023), b) Fe^2+^ never gets oxidized in the cytosol and proceeds into chloroplasts, or c) photoreduction of Fe^3+^, as shown previously (Bughio et al., 1997; Solti et al., 2012), plays a significant role in the supply of Fe^2+^ for chloroplast import. Import may occur via PIC1, which was not upregulated in *opt3*. Since *pic1* mutants are strongly compromised, *opt3pic1* alleles were not investigated. We did test the suggested Fe carrier MFL1 (Tarantino et al., 2011). Loss of MFL1 in the *opt3* background resulted in 20-25% reduced plastid Fe contents (fig. S3). This is in line with the observation of Tarantino et al. (2011) that *mfl1* only displays reduced leaf tissue iron levels from the Wt control if plants are fertilized with excess iron. Based on these observations, MFL1 might be a Fe transporter activated at Fe surplus conditions. Taken together, it is clear that the current understanding on plastid Fe transport needs further revisions.

This is especially urgent, since our study reveals the high significance for chloroplasts in sensing the plant’s general Fe status. When the plastid’s Fe storage capacity was decreased by eradicating stromal ferritins, Fe uptake in the absence of OPT3 ceased. Although IRT1 levels remained high in roots, substrate acidification mediated by AHA2 (Santi and Schmidt, 2009) and Fe reductase activity, catalyzed by FRO2 (Robinson et al., 1999), both declined. AHA2 (Santi and Schmidt, 2009) and FRO2 (Robinson and Downton, 1984) form a protein complex with IRT1 and are critical to facilitate Fe^2+^ uptake into plants (Martín-Barranco et al., 2020). Hence, our data suggest that Fe import in *opt3fer* mutants is dampened by reducing AHA2 and FRO2 activity. In addition, WAKL4 expression in *opt3fer* was strikingly upregulated. WAKL4 initiates NRAMP1 degradation by phosphorylation, effectively decreasing metal uptake via the carrier (Yuan et al., 2024). Although we found WAKL4 upregulation in leaves, reduction in NRAMP1 may contribute to attenuated Fe influx into *opt3fer* plants. Alternatively, root to shoot translocation of Fe could also be impaired.

*opt3fer* mutants grew slower than *opt3* plants. Since photosynthesis was unaffected, growth may be impacted from activated emergency programs once the capacity to safely store excess Fe in the ferritins is gone. Indeed, we found more DEGs in *opt3fer* compared to controls. Most dramatically enriched were GO terms related to Fe transport and homeostasis. DEGs within this cluster, basically the leaf ferrome genes (McInturf et al., 2021), were mostly downregulated. Hence, transcriptomics is in line with the observed decline of leaf and plastid Fe. In addition, we found many GO terms related to plant stress and defense. Here, the picture is more diverse. Loss of OPT3 and concomitant activation of Fe-starvation are known to confer broad pathogen resistance e.g., against necrotrophic fungus *Botrytis cinerea* (Trapet et al., 2020). Fe-starvation signaling initiates hormonal responses whereby JA and salicylic acid (SA) become antagonistically regulated via nitric oxide (NO) and ethylene (ET) to modulate the response (Herlihy et al., 2020). JA and Auxin-related DEGs were down-while ARACIN1 was upregulated in *opt3fer*. Indeed, ectopic ARACIN1 expression results in small, slow-growing plants (Neukermans et al., 2015), which likely contributes to the stunted phenotype *opt3fer*. Alternatively, the limited capacity to deposit Fe into plastids may activate ferroptosis. Herein, cells deplete GSH, accumulate ROS, and exhibit Fe-dependent lipid peroxidation, which in conjunction can trigger cell death (Dangol et al., 2019; Distéfano et al., 2020). In tobacco it has been shown that silencing the Fe storage protein ferritin triggers ferroptosis (Macharia et al., 2020). Hence, we posit that *opt3fer* grows slowly due to activated pathogen defense factors, ferroptosis or a combination thereof.

To track the subcellular fate of leaf Fe and other metals, we employed NAF and built a model of the physiological processes at work in our mutant panel (fig. 7). Fe data collected from wild-type Arabidopsis showing 70% of cellular Fe in the plastid fraction match historic data from sugar beets (79%) very well (Terry and Low, 1982). Overall, this suggests that the subcellular metal ion homeostasis may be conserved among glycophytic dicots. Our study expands the knowledge foundation by providing additional information on cytosolic and vacuolar levels for Fe but also several other nutrient metals.

**Figure 7.**
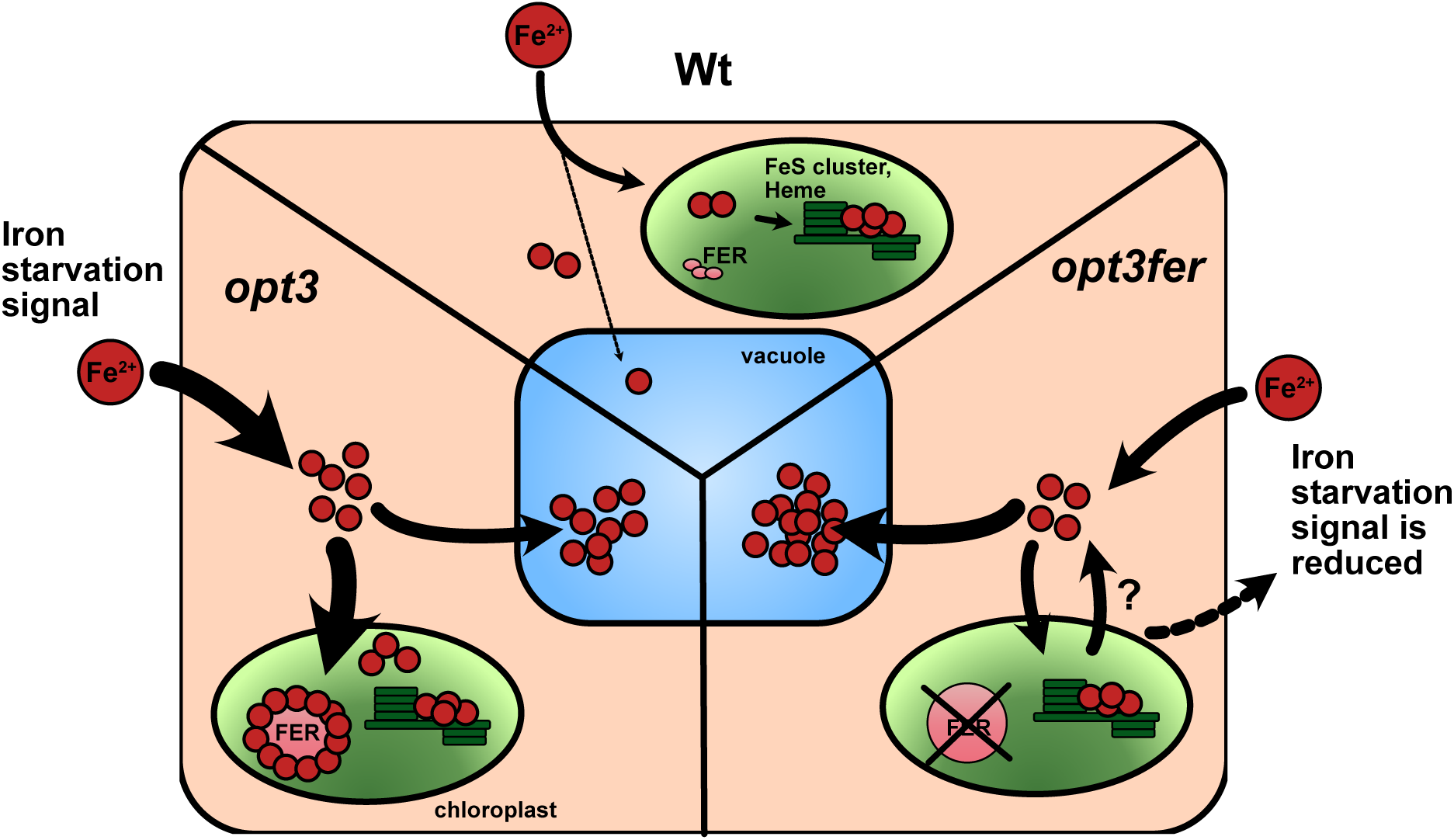
Working model. In Wt leaf cells chloroplasts represent the main Fe sink. Minor amounts of Fe are present in the cytosol or stored in the vacuole. In *opt3* mutants, Fe is over accumulating in the leaf cell due to a constitutive Fe starvation signal. Surplus Fe is stored in plastid ferritins and the vacuole. Upon loss of plastid ferritins (*opt3fer*), Fe is mainly stored in the vacuole. However, the vacuole is not able to compensate the Fe storage capacity of plastid ferritins. As a result, a signal is released which suppresses the constitutive Fe starvation signal in *opt3* and reduces Fe uptake into the root.

In the absence of OPT3 and ferritins, we found that the central vacuole becomes a backup Fe storage. Although *opt3* mutants already sequester twice as much Fe into the vacuole than into plastids, relatively speaking *opt3fer* pumps even more Fe into vacuoles, since the mutant’s total leaf Fe content drops by almost half compared to *opt3*. *opt3fer* may also utilizes the apoplast to decrease Fe toxicity or other compartments we could not probe due to technical limitations. With approximately 10 - 30% of total leaf Fe stored as apoplastic Fe (apoFe), apoFe represents a significant Fe storage (Liu et al., 2023). However, the decrease in PDF transcripts, which encode apoFe-binding proteins, renders this option less likely.

Lastly, we utilized the mutant set to investigate how plants deprived of plastid ferritins cope with Fe starvation. Fe release from plastid ferritins akin to the system in mammals has only been investigated *in vitro* thus far (Laulhere and Briat, 1993). Wt plants transferred into Fe starvation conditions exhibit chlorosis and low *F*_v_/*F*_m_ in old leaves. This effect was mostly independent of the presence of ferritins and occurred at both Fe pre-starvation concentrations (25 vs 100 μM). In contrast, *OPT3*-deficient plants, which cannot remobilize Fe, maintained much higher *F*_v_/*F*_m_ in old tissue and did not display leaf chlorosis. Therefore, young leaves in *opt3* and *opt3fer* were more affected than Wt and *fer* mutant controls. Again, starvation effects were independent of the presence of ferritins. Altogether our experiments show that A) Arabidopsis does remobilize Fe to a substantial degree; B) absence of ferritins *in vivo* had little effect on Fe remobilization. This suggests that plant ferritins, akin to the structurally more similar bacterial ferritins (Wade et al., 1993), function primarily to prevent Fe from reacting with oxygen and to trigger oxidative stress (Briat et al., 2010). Since *fer* also displayed enhanced tolerance to iron starvation if grown at iron excess conditions, alternative iron storages like the vacuole must be present. Ferritin degradation and secure Fe release is probably realized by vacuolar autophagy late in the vegetative stage i.e., during leaf senescence.

In conclusion, this study successfully describes the chloroplast ionome of several glycophytic dicots, including the model species *A. thaliana*. Among these species, plastid metal contents were highly similar, suggesting that photosynthesis and other critical pathways inside the organelle function within a fairly small range of nutrient ions and buffer conditions. Natural metal hyperaccumulator species have adopted mechanisms to protect the plastid ion homeostasis, which will be of great interest to unveil. Nevertheless, it is possible to genetically manipulate the chloroplast ionome and for instance force it into becoming a site of strong Fe accumulation. Fe biofortification of plant-based diets is of great interest. Since plastids and ferritins can be readily and cost-effectively purified, our study provides new avenues into utilizing leaf ferritins as biotechnological targets to supplement vegetarian food with Fe.

## Material and Methods

### Plant growth and isolation of *Arabidopsis* mutant lines

Plants (*A. thaliana* ecotype Col-0 and *P. sativum* ecotype sparkle) were cultivated on soil (Substrate A210, Stender, Schermbeck, Germany) at long day conditions (16 h illumination / 8 h dark) at 110 µmol photons m ^-2^ s ^-1^, 22°C, 50% humidity. Plants were harvested at 21-days for all experiments if not stated otherwise. For chloroplast isolation plants were kept in the darkness for 24 hours pre-isolation to lower the starch content. *Arabidopsis halleri* ssp. *halleri* (O’Kane and Al-Shehbaz, 1997) population Langelsheim, accession Lan3.1, (Becher et al., 2004) was cultivated on greenhouse soil (Minitray soil, Balster Einheitserdewerk, Frödenberg) trough vegetative propagation and harvested at RUB, Bochum. T-DNA mutants were ordered from NASC. Homozygous genotypes were confirmed by PCR using primer combinations depicted in tab. S1. Higher order mutants were obtained by crossing the respective single and higher order mutant alleles. Genotypes were confirmed by PCR using the primer combinations depicted in tab. S1.

To obtain *A. thaliana* mutant roots for immunodetection of IRT1 and FCR assays, plants were cultivated vertically on 1/2-Murashige & Skoog (1/2-MS) media (Murashige and Skoog, 1962).

### Plant cultivation in hydroponic media for Fe starvation experiments

For Fe starvation 1/10-Hoagland media was prepared according to Becher et al. (2004) with minor adjustments using the following concentrations (1.5 mM Ca(NO_3_)_2_, 0.28 mM KH_2_PO_4_, 0.75 mM MgSO_4_, 1.25 mM KNO_3_, 0.5 µM CuSO_4_, 5 µM ZnSO_4_, 5 µM MnSO_4_, 25 µM H_3_BO_3_, 0.1 µM Na_2_MoO_4_, 50 µM KCl, 3 mM MES-KOH pH 5.7, without (0), 25 µM or 100 µM Fe-EDDHA as indicated, 0.6% (w/v) agarose). Seeds were sterilized using 70% (v/v) EtOH. Plants were cultivated in sterile conditions at long day conditions (16 h illumination / 8 h dark) at 110 µmol photons m ^-2^ s ^-1^. After 10 days growth at control conditions (0, 25, or 100 µM Fe-EDDHA) seedlings were shifted either to plates containing 0 µM Fe-EDDHA for Fe starvation or to plates with the same Fe-EDDHA concentrations as a control. All plants were subjected to PAM-analysis and subsequent harvest at 14 days post shifting.

### Stable transformation of *Arabidopsis thaliana*

*FRO7*, *FRO6* and *FER1* genomic DNA (ATG – STOP) was amplified by PCR using the primers given in tab. S1. Due to the sequence similarity between *FRO6* and *FRO7*, *FRO6* gDNA was first amplified including the non-translated region. *FRO6* genomic DNA was subsequently amplified from this intermediary template. All constructs were cloned using the pGreen 2.0 system using the UBQ10 promotor and a HSP18.2 terminator (Pratt et al., 2020) via Gibson cloning. For subsequent detection all constructs were fused with a tobacco etch virus restriction enzyme (TEV) - mVenus tag. *A. thaliana* Col-0 were transformed via floral dip (Clough and Bent, 1998) using the GV3101 agrobacteria strain outfitted with help plasmid pSOUP (Hellens et al., 2000). Transgenic plants were identified on ½ MS plants containing 25µg/ml (w/v) hygromycin. Microscopy was carried out in the T_1_ generation. Immunoblotting in fluorescence-selected F_2_ plants.

### Chloroplast isolation

Intact chloroplasts were isolated according to Aronsson and Jarvis (2002). In brief: 21-day old plants were incubated in darkness for 24 h to remove internal starch. Next, plants were harvested and carefully broken down in pre-cooled isolation buffer (300 mM sorbitol, 5 mM MgCl_2_, 5 mM EDTA, 20 mM HEPES / NaOH pH 8.0, 50 mM ascorbic acid, 0.1% (w/v) BSA) using a Polytron homogenizer (Bachhofer, Reutlingen, Germany). Subsequently, the suspension was filtered through gauze and centrifuged at 1500*g*, 3 min, 4°C. Total chloroplasts were resuspended in chloroplast isolation buffer and separated on an 82% (v/v) / 30% (v/v) 2-step percoll gradient (82% (v/v) / 30% (v/v) percoll (GE-healthcare, Chicago, USA), 300 mM sorbitol, 5 mM MgCl_2_, 5 mM EDTA, 20 mM HEPES / NaOH pH 8.0). Separation of chloroplasts into “intact chloroplasts” and “broken chloroplasts” was obtained by centrifugation of the gradient for 10 min at 2000*g*, 4°C. Subsequently, intact chloroplasts were resuspended in chloroplast washing buffer (300 mM sorbitol, 5 mM MgCl_2_, 5 mM EDTA, 20 mM HEPES / NaOH pH 8.0) and centrifuged at 1000*g*, 5 min, 4°C. For element determination isolated chloroplasts were resuspended in chloroplast washing buffer to a chlorophyll concentration of 0.4 µg chlorophyll / µl. Chlorophyll concentration was determined in 80% (v/v) acetone by measuring the absorption at 663, 645 and 750 nm (Arnon, 1949). Chlorophyll content was calculated using the following equation:

µg chlorophyll / µl sample= 8.02 x (645 nm – 750 nm) + 20.2 x (663 nm-750 nm)

For every chloroplast sample a fraction of the corresponding leaf tissue material was kept for leaf tissue element determination.

### Element determination

Chloroplast and leaf tissue elements were analyzed by total X-ray fluorescence (TXRF) using a S4 T-STAR (Bruker, Berlin, Germany). Isolated chloroplasts were mixed with element standards in a 1:1 ratio to a final concentration of 50 ppm Scandium (WL) and 1 ppm Gallium (Mo-K), 0.1 % (w/v) polyvinyl alcohol (PVA). For background determination, a sample of chloroplast washing buffer was measured. Leaf tissue was dried at 60°C and homogenized in a zirconium mortar. Three mg of leaf powder was digested in 200µl 69% (v/v) HNO_3_ at 95°C for 90 min. Digested samples were mixed with element standards in a 1:10 ratio to a final concentration of 50 ppm Scandium (WL) and 1 ppm Gallium (Mo-K). For background determination, a sample of 69% (v/v) HNO_3_ was measured (Höhner et al., 2016). All samples were spotted on silicon coated quartz glass carriers and measured at 50 kV, for 1000 sec. (isolated chloroplasts) or 600 sec. (leaf tissue) per sample using Mo-K and WL excitation. Subsequently, the element concentration was normalized on the chlorophyll concentration or dry weight of the respective sample. Background element content (elements present in chloroplast washing buffer or 69% (v/v) HNO_3_ for leaf tissue elements) were subtracted from the respective element measurement. Significant differences (*p* < 0.05) between samples were determined using an ordinary one-way analysis of variance (ANOVA) with Tukeýs multiple comparison test using GraphPad Prism 10.4.1 (Dotmatics, Boston, USA).

### Leaf chlorophyll determination

Leaf chlorophyll levels were determined according to Lichtenthaler (1987), Porra et al. (1989). In brief, 20 mg of nitrogen ground leaf tissue material was weighed in a pre-cooled reaction tube. Chlorophyll was extracted using 80% acetone overnight. Debris was removed by centrifugation at 10.000*g* for 5 min at 4 °C. Absorption of the supernatant was measured at 646 nm and 663 nm. Chlorophyll content normalized on fresh weight was calculated using the following equation:

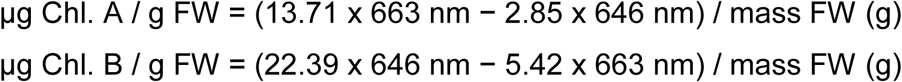

### Immunoblotting

Immunoblotting was carried out either on protein isolates acquired from isolated chloroplasts or total leaf tissue (Völkner et al., 2021; Völkner et al., 2024). In case of chloroplasts, isolated chloroplasts were pelleted and resuspended in 1x SDS-loading dye (1% (v/v) glycerol, 5% (v/v) β-mercaptoethanol, 2% (w/v) SDS, 62.5 mM Tris-HCl pH 6.8, 0.003% (w/v) bromophenol blue) and heated at 95°C for 5 minutes. In case of leaf tissue, equal amounts of leaf tissue were ground and solubilized in solubilization buffer (10% (v/v) glycerol, 150 mM NaCl, 2 mM EDTA, 5 mM DTT, 50 mM Tris-HCl pH 7.5, 1% (w/v) SDS, plant protease inhibitor mix) for 30 min at 4°C. Subsequently the isolates were mixed with SDS-loading dye and boiled at 95°C for 5 min. Proteins were separated on a 10.5% (w/v) - 15% (w/v) SDS-PAGE depending on the molecular weight of the protein of interest and transferred to a PVDF membrane using a wet-blotting system. Immunodetection was carried out with the respective antibody at 4°C overnight (FER1: AS152898, IRT1: AS111780, CSD2: AS06170, FeSOD: AS06125, obtained from Agrisera, Vännäs, Sweden, PIC1: (Duy et al., 2007)). Target proteins were visualized using enhanced chemiluminescence by secondary antibodies coupled to horse radish peroxidase.

### Photosynthetic parameters

For determination of photosynthetic performance, plants were analyzed using a Walz Imaging PAM (Walz GmbH, Effeltrich, Germany) as described in Kunz et al. (2009). In brief, plants were dark-incubated for 30 min prior measurement. Photosynthetic performance was measured using a standard induction curve at 110 µmol photons m^−2^ s^−1^ for 300 sec in 20 sec steps. Analysis and export of false color images was carried out using the Walz ImagingWinGigE software.

### Imaging of isolated *Arabidopsis* protoplasts

Protoplasts were isolated as described by Wu et al. (2009). Imaging was performed using a Stellaris 5 Laser Scanning Confocal Microscope (Leica, Wetzlar), equipped with a 405 nm diode and a supercontinuum Wight Light Laser (WLL). Emission was detected with a Power Hybrid HyD S detector. For protein localization, recombinant proteins fused to mVenus (YFP) were excited at a wavelength of 514 nm, and emission detected at 520-580 nm. Chlorophyll autofluorescence was excited at 405 nm and emission recorded at 623-813 nm. Protoplasts were imaged acquiring a Z-stack, enhanced by LIGHTNING as required, followed by processing using the LAS X software to generate maximum intensity projections.

### Blue Native (BN) PAGE

BN-PAGE on isolated chloroplasts was carried out as described by (Nickel et al., 2016; Völkner et al., 2021; Völkner et al., 2024) with minor modifications. In brief, isolated chloroplasts equal to 15 µg chlorophyll per sample were solubilized for 15 minutes on ice using 0.5% (w/v) dodecyl maltoside. After solubilization, samples were mixed with BN-loading dye (750 mM aminocaproic acid, 5% (w/v) Coomassie G-250) and separated on a 5 – 15% (v/v) native acrylamide gradient.

### RNA isolation and sequencing

Total RNA was isolated from 21-day old leaf-rosettes using the Macherey-Nagel plant RNA isolation kit (Macherey-Nagel, Düren, Germany) according to the manufacturer’s instructions including on-column DNAse digest. Total RNA was sequenced by BMKGene (Münster, Germany) with Illumina 150bp paired end sequencing.

### Analysis of RNA-SEQ data

Analysis of RNAseq data was carried out similar to DeTar et al. (2021). Raw sequencing data were deposited to NCBI, SRA (PRJNA1216402). Obtained reads were processed using a pipeline on the Galaxy platform (Community, 2024) in including the tools TrimGalore!, RNAstar, feature counts, limma-voom, venn diagram, and heatmap2 with default settings. Mapping was carried out to the reference genome obtained from Tair10 (Berardini et al., 2015). Differentially expressed genes (DEGs) with an adj. P. value < 0.05 were identified as significant DEGs. For general overview on significant DEGs a cutoff of log_2_FC > 0.1 for upregulated or < -0.1 for downregulated genes was applied. For gene ontology (GO) term analysis, a cutoff of log_2_FC > 1 or < -1 (adj. P. value < 0.05) was applied. Gene ontology terms were analyzed by g:Profiler (Kolberg et al., 2023) and visualized using revigo (Supek et al., 2011) and the cytoscape software. Ferrome genes were identified according to McInturf et al. (2021).

### Non-aqueous fractionation

Non-aqueous fractionation was carried out on 21-day old seedlings as described by (Hernandez et al., 2023). Plant material was quenched in liquid nitrogen, ground to a fine powder and lyophilized. The lyophilized powder was then suspended in a mixture of n-heptane and tetrachlorethylene (nH-TCE), sonicated and centrifuged at 4°C with 20,000g. The supernatant was transferred to a new tube and the pellet was suspended in a nH-TCE solution with increased density. This procedure was repeated until a maximal density of 1.6 g cm^-3^ (i.e., pure TCE). The pellets were suspended in n-heptane and split into subfractions (i) marker enzyme activity measurements, and (i) element analysis. The fractions were dried in a desiccator before further use. Marker enzyme activities were determined for plastids (alkaline pyrophosphatase), cytosol (UDP-glucose pyrophosphorylase) and vacuole (acidic phosphatase). Distribution of marker enzymes in the respective fractions was depicted as geom smoothed curves using R-studio.

Lyophilized samples were extracted in 100 µl 69% (v/v) HNO3 for 90 minutes at 95°C. Extracted samples were mixed 1:10 with element standards to a final concentration of 1 ppm gallium and 50 ppm scandium. Samples were spotted on silicone coated quartz glass carriers and measured at 50 kV for 1000 s using TXRF (S4 T-STAR, Bruker, Berlin, Germany) MoK and WL excitation.

Subcellular distributions of elements were determined using the NAFalyzer app (https://github.com/cellbiomaths/NAFalyzer).

### Determination of root ferric chelate reductase activity

Activity of Fe(III) reductase activity was determined as described by Grillet et al. (2018). Roots were harvested from plants were cultivated on 1/2-MS for 2 weeks. Next, the roots were incubated at gentle shaking in the dark in 1 ml essay solution (100 μM Fe^3+^-EDTA and 300 μM bathophenanthroline disulfonate (BPDS), 10 mM MES pH 5.5). After 1 h the Fe^2+^BPDS_3_ concentration and therefore the activity of the root FCR was determined by measuring the absorbance at 535 nm and normalized on root fresh weight. Significant differences were analyzed by one-way ANOVA.

### Determination of root rhizosphere acidification

To determine the activity of AHA2 at the root, rhizosphere acidification was measured as described previously (Agrahari et al., 2024). In brief, 1-week old seedlings precultured on ½-MS were shifted pH indicator plates (92.6 µM bromocresol purple, 9 mM CaCl_2_, 13.4 mM KCl, 7.5% (w/v) plant agar). Plants were incubated for 2-4 h until the change in pH at the rhizosphere was visible.

### Enzymatic activity of stromal NADP malate dehydrogenase

Isolated chloroplasts were adjusted to a concentration of 0.6 µg chlorophyll * µl^-1^ for all samples and mixed with DTT to a final concentration of 20 mM DTT. Next up, each chloroplast sample was split in two samples of the same volume (500µl) of which one sample was sonicated to lyse the chloroplasts. The other sample was not treated. Subsequently, 10 µl of chloroplast suspension of both samples was mixed with reaction buffer (300 mM sorbitol, 5 mM MgCl_2_, 250 µM NADPH, 20 mM HEPES-KOH pH 8.0) in a 96 well plate. Next, oxaloacetate was added simultaneously to all samples. Activity of the stromal NADP-malate dehydrogenase was determined by measuring the absorption of NADPH at 340 nm for 30 min in 30 sec steps in a Spark plate reader (Tecan, Männedorf, Switzerland). Data collection was started after 5 min post reaction start to settle the reaction. Subsequently, the first NADPH absorption measurement for both sonicated and non-treated chloroplasts was set to 1 for visualization. NADP-MDH activity was depicted as the inverted change of NADPH absorption rate.

### Alignment of protein sequences

Protein sequences for alignment were accessed from Tair. Subsequently, alignment of the sequences was performed using Clustal Ω (Madeira et al., 2024). Visualization of the alignment was carried out using Jalview (Waterhouse et al., 2009).

### Accession numbers

*FER1* (AT5G01600), *FER3* (AT3G56090), *FER4* (AT2G40300), *FRO7* (At5g49740), *FRO6* (At5g49730), *OPT3* (AT4G16370), *MFL1* (At5g42130), *IRT1* (AT4G19690) *opt3-2* (SALK_021168C), *opt3-3* (SALK_058794C), *fer1-1* (SALK_055487), *fer3-1*(GABI-KAT_496A08), *fer4-1*(SALK_068629), *fro7-1*(Salk_027287), *fro7-2* (Salk_048230), *mfl1-1* (SALK_056579), *mfl1-2* (SALK_007617)*, irt1-2* (SALK_054554)

### Author Contributions

- H-H.K. designed research and wrote the manuscript. L.J.H. cloned constructs, isolated plant mutants, performed most experiments, analyzed data, and wrote the manuscript. L.Ö F. isolated mutants, performed element analysis and conducted final Fe starvation assays. S.M. performed microscopy and designed cartoons. A.M. established the initial experiments on Fe starvation, cloned constructs and generated transgenic lines for microscopy. T.N. and L.S. prepared NAF samples and analyzed NAF data. R.A.D. performed initial plastid element analyses. K.P. provided the PIC1 antibody. D.M.C. provided the *opt3* germplasm. U.K. provided *A. halleri* plants and performed confirmatory multi-element analyses by Inductively-Coupled Plasma Optical Emission Spectrometry (ICP-OES). B.B. analyzed data and helped with writing and editing. All authors assisted in editing the manuscript.
- Funding: H-H.K., L.J.H, and S.M. were funded by the Deutsche Forschungsgemeinschaft (DFG) (SFB-TR 175, project B09). T.N. was supported by the Deutsche Forschungsgemeinschaft (DFG) (SFB-TR 175, project D03). Research by D.M.C. is supported by the US NSF award MCB-2224839. Early project funding came from an NSF Career Award (IOS-1553506) to H-H.K. Open Access funding enabled and organized by Projekt DEAL.

## Acknowledgements

We are grateful to Dr. Ricarda Höhner (previously at WSU) who performed first TXRF analyses on isolated plastids. We thank gardener Albert Schorer from LMU for his support and insights and Andreas Aufermann from RUB for *A. halleri* cultivation. Thanks to Dr. Stephane Mari (INRAE Institut des Sciences des Plantes Montpellier, France) for providing *fer1,3,4* seeds. We thank Drs. Janneke Balk (John Innes Centre, UK) and Javier Romera (University of Cordoba, Spain) for providing pea sparkle germplasm.

---

**Figure S1.**
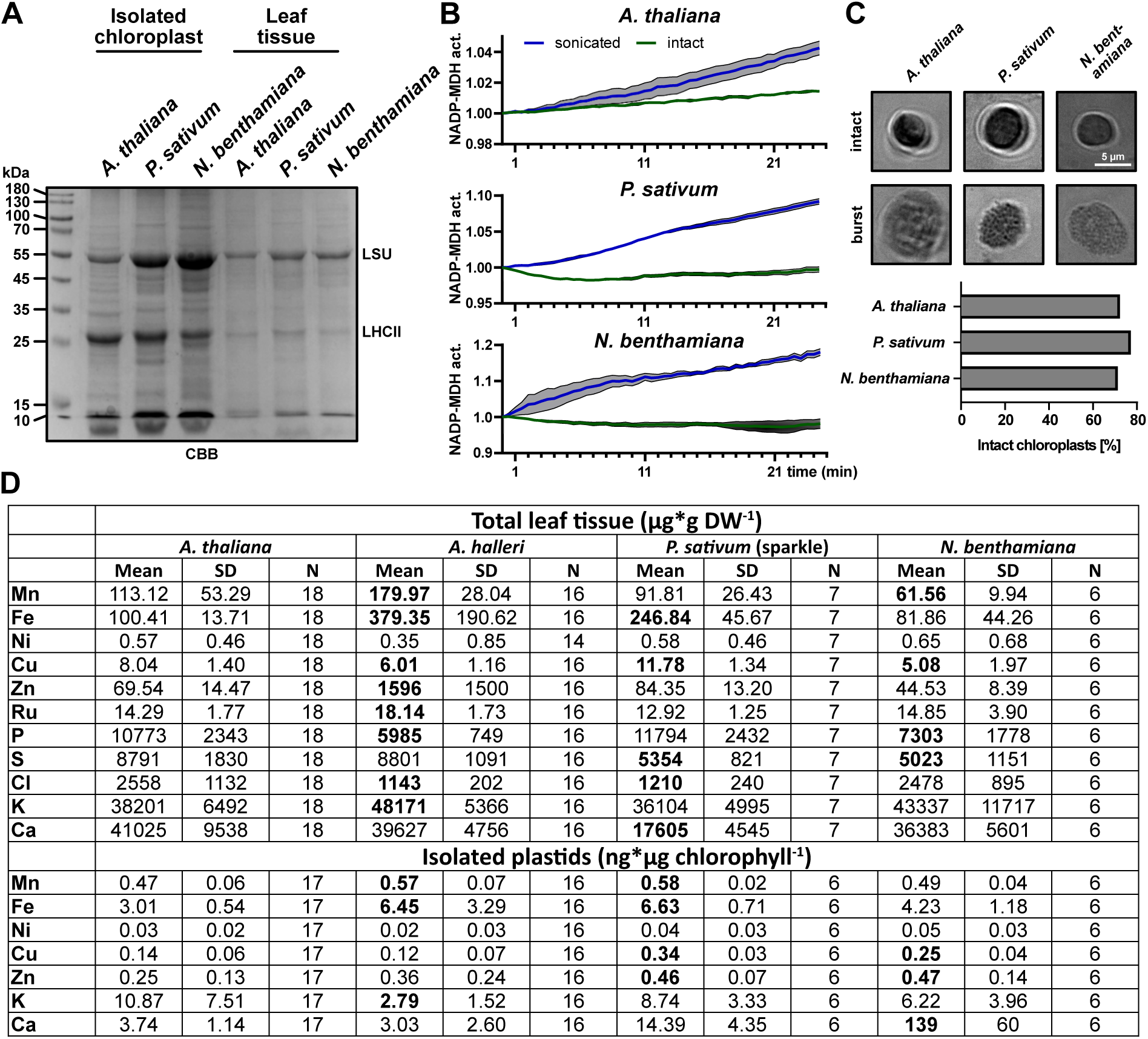
Leaf and plastid ionome of four plant species. (**A-C**) Quality control of plastids isolated from *A. thaliana*, *P. sativum* and *N. benthamiana* used in this study. (**A**) The presence of the stromal localized large subunit of RuBisCO (LSU) was probed using a Coomassie Brilliant Blue (CBB) stained SDS-PAGE. (**B**) The stability of isolated plastids was probed by determining the activity of the stromal localized NAD malate dehydrogenase (MDH) over a time period of 25 minutes. (**C**) Isolated chloroplasts were grouped by their visual appearance into “intact” or “burst” chloroplasts based on the presence of the characteristic white “halo” surrounding intact chloroplasts. (**D**) Full set of leaf tissue and plastid ionome of *A. thaliana*, *A. halleri*, *P. sativum* and *N. benthamiana*. Elements were determined by TXRF. Significant differences (stated in bold) in other plant species from *A. thaliana* within one element were obtained using one-way ANOVA (*p* < 0.05).

**Figure S2.**
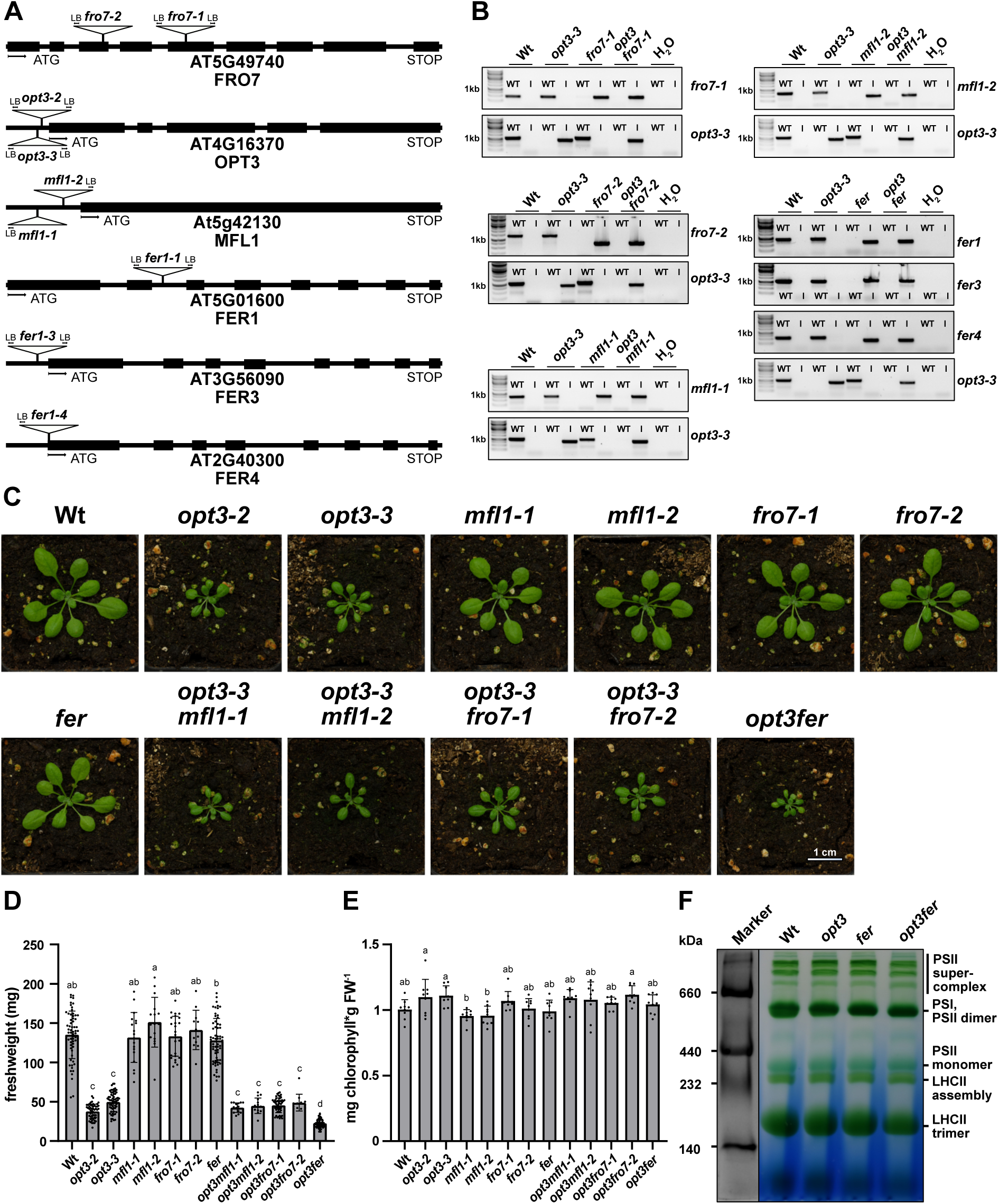
Characterization of plant lines used in this study. **(A)** Annotation of T-DNA insertion of lines used in this study. (**B**) Confirmation of single and higher order mutants by PCR. (**C**) Visual phenotype of mutant lines used in this study. (**D**) Freshweight of mutant lines used in this study. Data is presented as mean ± SD. Significant differences between lines were observed using one-way ANOVA (*p* < 0.05) indicated by different letters. (**E**) Leaf chlorophyll of mutant lines used in this study. Data is presented as mean ± SD. Significant differences between lines were observed using one-way ANOVA. (**F**) Analysis of the major photosynthetic complexes by Blue Native PAGE.

**Figure S3.**
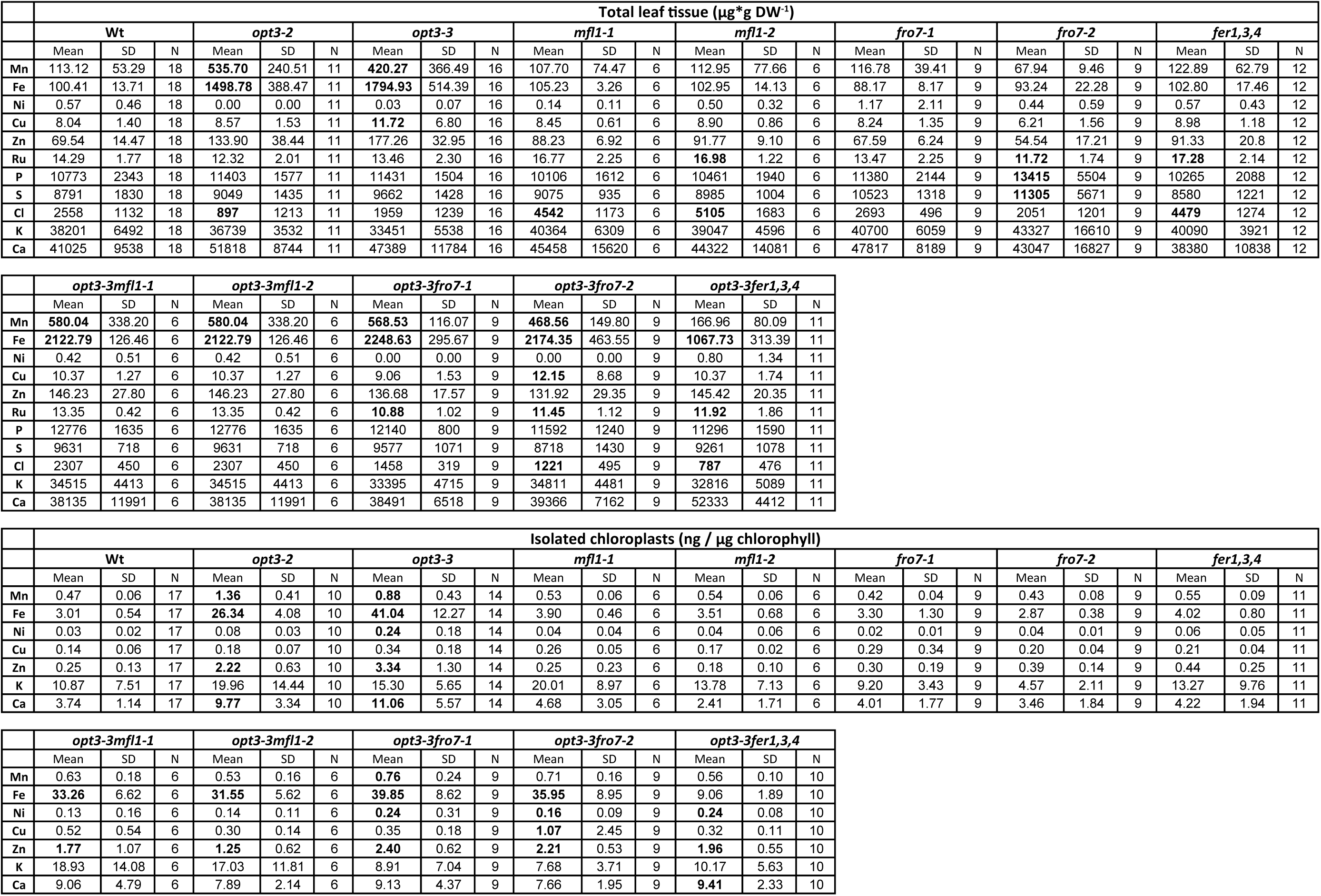
Leaf tissue and plastid ionome of all mutant lines used in this study. Elements were determined by TXRF. Leaf tissue element levels were normalized on dry weight, chloroplast element levels on chlorophyll respectively. Significant differences (stated in bold) in a mutant from the Wt within one element were identified using one-way ANOVA (*p* < 0.05).

**Figure S4.**
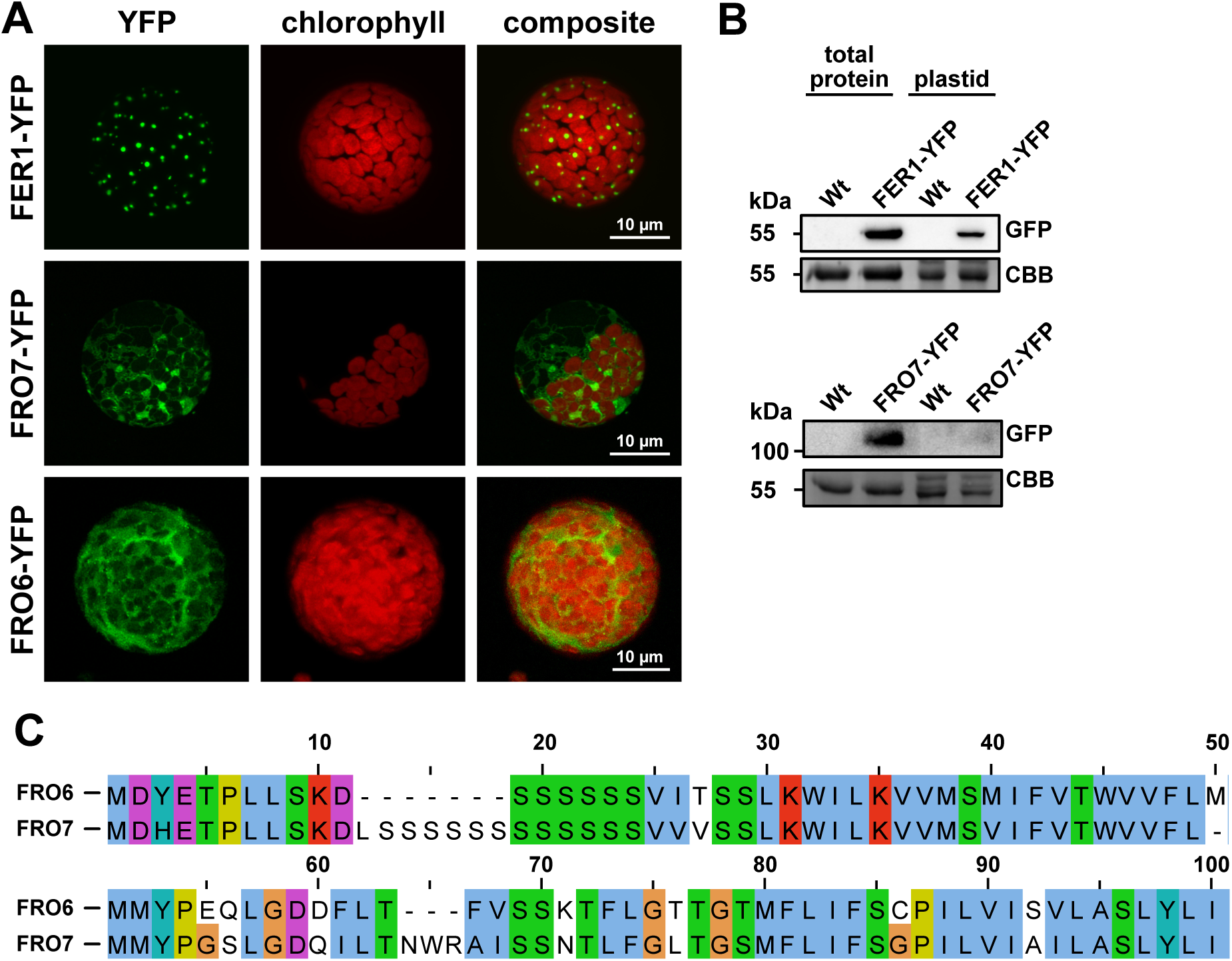
FRO7 is not localized in chloroplasts. **(A)** Localization of FRO6, FRO7 and FER1 was analyzed by fusing the respective protein with a C-terminal fluorescent mVenus (YFP) tag. Data is presented as YFP channel, chlorophyll autofluorescence and merge. (**B**) Localization of the YFP-tagged protein to plastids was analyzed by performing a immunoblot using an GFP antibody. (**C**) Alignment of the 100 N-terminal residues of FRO6 and FRO7.

**Figure S5.**
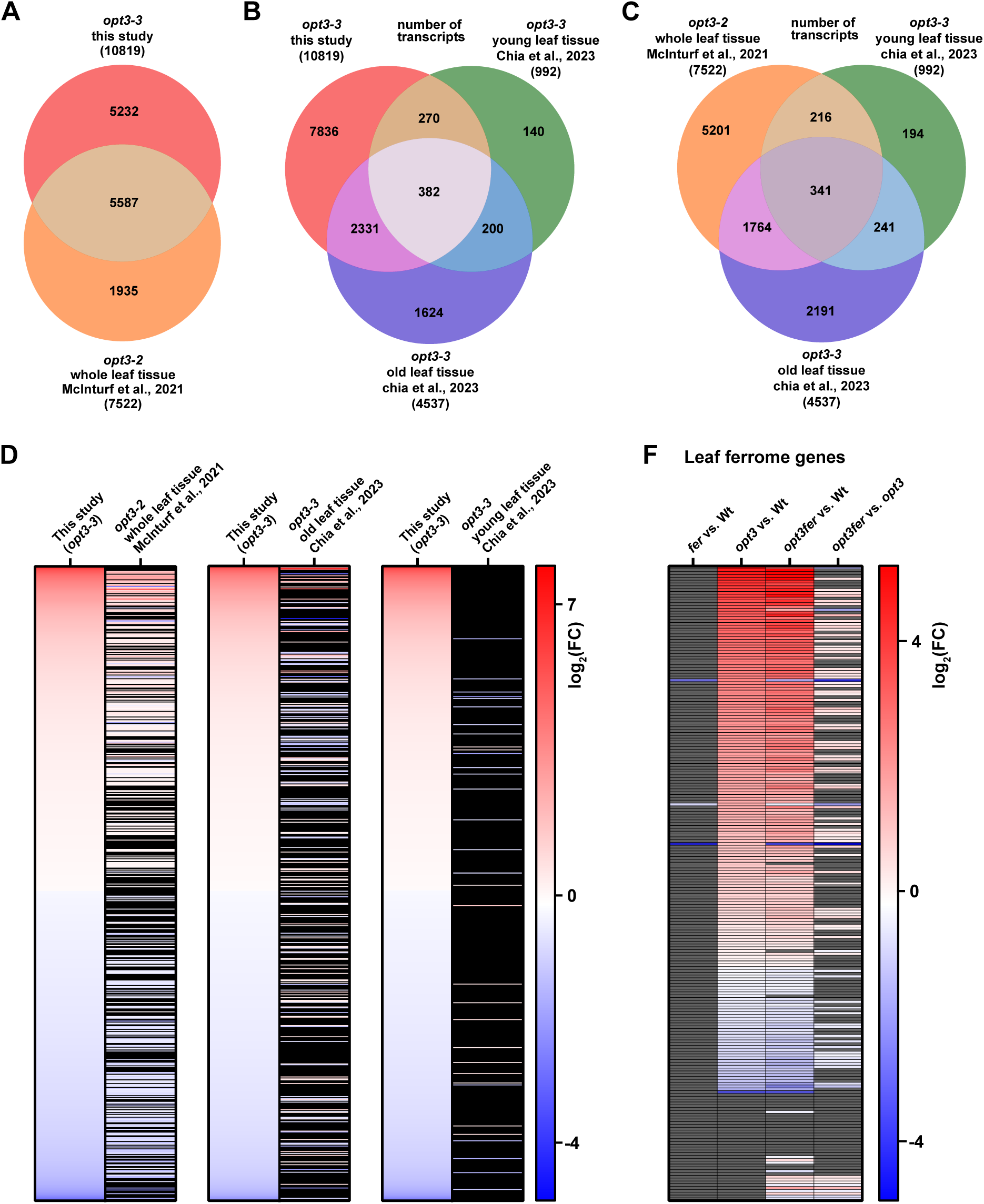
Comparison of DEGs identified in this study with previous studies. (**A-C**) Comparison of significant DEGs (log_2_FC **≥** 0.1, *p* < 0.05) identified in this study with previous studies (McInturf et al., 2021; Chia et al., 2023). Venn diagrams were obtained using the galaxy platform. (**D**) Detailed comparison on significant DEGs (log_2_FC **≥** 0.1, *p* < 0.05) identified in this study with previous studies. We identified a high level of similarity between the DEGs identified in our study and DEGs identified in *opt3-2* (McInturf et al., 2021). We identified less similarity with the DEGs identified by Chia et al. (2023) most likely due to the separation in young and old leaves in the Chia et al. (2023) study. (**E**) Significant DEGs in this study (log_2_FC **≥** 0.1, *p* < 0.05) filtered for ferrome genes proposed by McInturf et al. (2021). Non-significantly changed DEGs are colored in gray.

**Figure S6.**
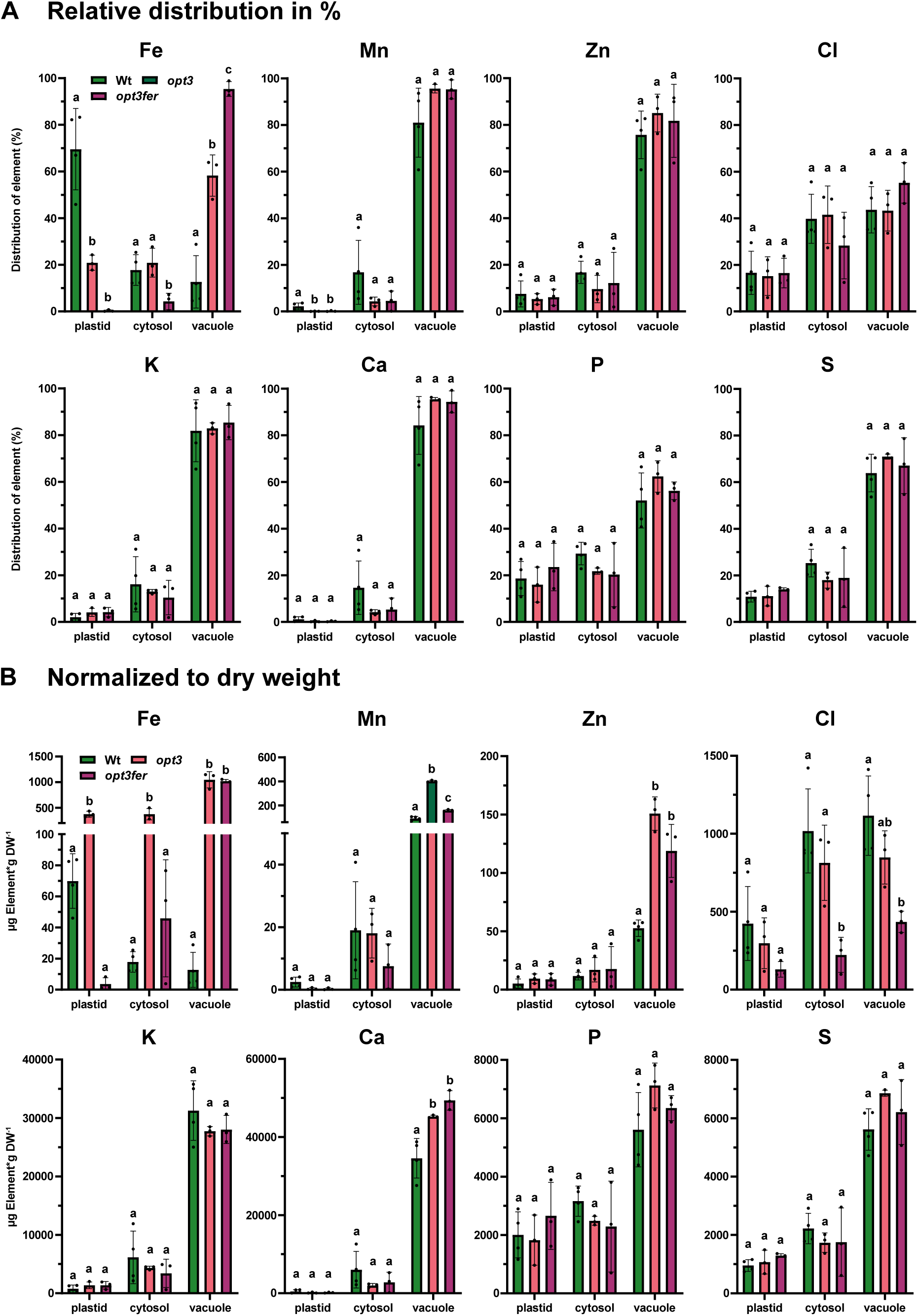
Subcellular distribution of elements identified by non-aqueous fractionation (NAF). **(A)** Relative distribution of elements detected via TXRF in organelles identified by NAF. Data is presented as mean ± SD. Significant differences between Wt, *opt3* and *opt3fer* were identified using one-way ANOVA (*p* < 0.05) indicated by different letters. (**B**) Relative distribution displayed in (A) was multiplied with the mean dry weight level of the respective element to obtain element levels within one organelle of the respective mutant. Data is presented as mean ± SD. Significant differences between Wt, *opt3* and *opt3fer* were identified using one-way ANOVA (*p* < 0.05) indicated by different letters.

**Figure S7.**
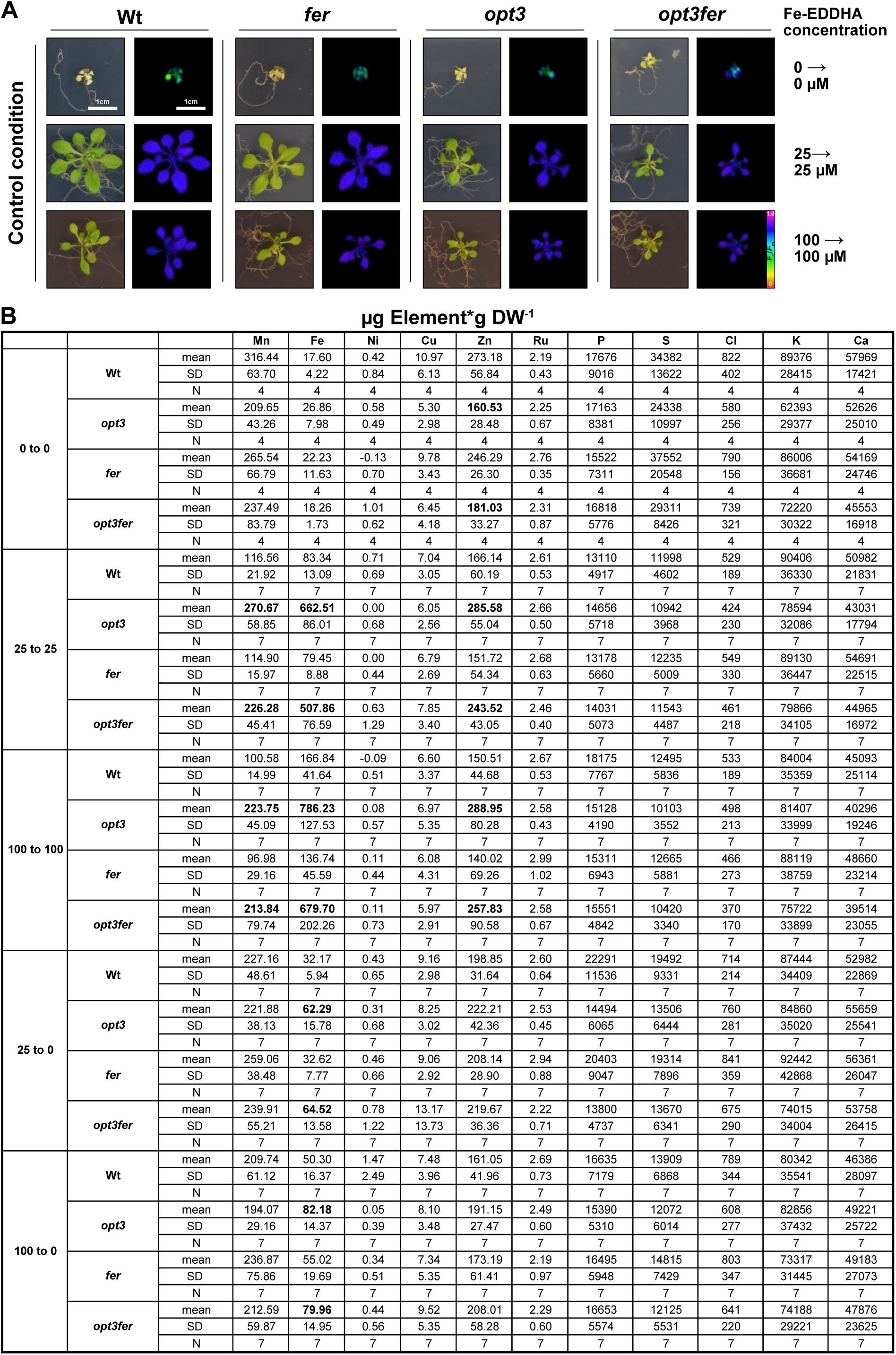
Elements of mutant lines at control and Fe starvation conditions. **(A)** Visual phenotype and photosynthetic performance (*F*_v_/*F*_m_) of Wt, *fer*, *opt3*, and *opt3fer* at control conditions. (**B**) All elements obtained via TXRF for lines cultivated on control conditions and shifted from control conditions to Fe starvation. Significant differences (stated in bold) from the were observed by one-way ANOVA (*p* < 0.05) within one treatment and one element.

